# Heterchronic shifts in a timing-keeping microRNA are associated with multiple instances of neoteny in plants

**DOI:** 10.1101/2025.04.30.651569

**Authors:** Aaron R. Leichty, R. Scott Poethig

## Abstract

The prolonged production of juvenile traits and an associated reduction or loss of adult traits during development (neoteny) can either arise from a change in genes that mediate the timing of the shift between juvenile and adult traits (timing genes) or from changes in genes that are necessary for the development of adult traits (response genes). To date, the relative contribution of each developmental mechanism to the origins of neoteny remain unclear. We examined this question in the plant genus, *Acacia*, which contains species that undergo the juvenile-to-adult vegetative transition (vegetative phase change) early in shoot development, as well as species that remain permanently juvenile, or have delayed vegetative phase change. Mapping the timing of vegetative phase change onto a molecular phylogeny of *Acacia* revealed that permanent juvenility has evolved multiple times and is sometimes associated with a delay in vegetative phase change in related species. In three cases, the absence or delay in vegetative phase change was associated with either higher amounts or a delayed decline in level of miR156, the master regulator of vegetative phase change in plants. These findings support the hypothesis that neoteny in *Acacia* has evolved not by a loss in the capacity to produce the adult leaf phenotype, but by a change in the timing of genes that promote juvenile leaf identity.

**SIGNIFICANCE STATEMENT:** Natural variation in the timing of the juvenile-to-adult transition has been described in many plant species, but the mechanism of this variation and its contribution to plant evolution is unknown. The genus *Acacia* is an excellent system in which to study these questions because it includes species that produce juvenile and adult leaves, as well as species that only produce juvenile leaves. Our results suggest that many permanently juvenile leaf species in *Acacia* are the result of the prolonged expression of the juvenile vegetative phase, not a loss of the ability to produce an adult leaf. In at least some cases, this neotenous phenotype is associated with a change in the expression of miR156, the master regulator of vegetative phase change.

## INTRODUCTION

Paedomophosis—the retention of juvenile features into sexual maturity—is one of the best known patterns of heterochrony in organismal evolution (Gould 1977). This phenomenon can arise either from the acceleration of reproductive maturation (progenesis), or from a delay in the transition between juvenile and adult phases of somatic/vegetative development (neoteny). In both cases, the mechanism(s) of this evolutionary shift in morphology remain largely unknown (Keyte and Smith 2014). This study concerns the mechanism of neoteny in *Acacia*, a genus of trees that contains species with well-defined juvenile and adult vegetative phases, as well as species that only produce juvenile vegetative traits. In principle, these neotenous species could arise in either of two ways. One possibility is a change in the activity of genes that regulate timing of the developmental transition (i.e. heterochronic or timing genes (Smith 2003; Moss 2007; Geuten and Coenen 2013), resulting in a failure or a delay in this transition (Figure 1A, middle route). A second possibility is that the genes responsible for executing the adult phenotype (response genes) become inactive; in this case, the timing mechanism remains intact, but the plant is incapable or attenuated in its ability to produce adult traits (Figure 1A, bottom route) (Roodt et al. 2019; Onyenedum and Pace 2021). One way to assess the likelihood of these scenarios is to compare the timing of developmental transitions in closely-related species. If species that display a delay in the juvenile-to-adult transition are phylogenetically close to permanently juvenile species, this would suggest that the absence of adult traits in these latter species is the result of a change in developmental timing, not a loss of the potential to produce adult traits. Additional evidence for a change in developmental timing would be provided by a change in the expression of genes known to regulate this developmental transition.

**Figure 1.**
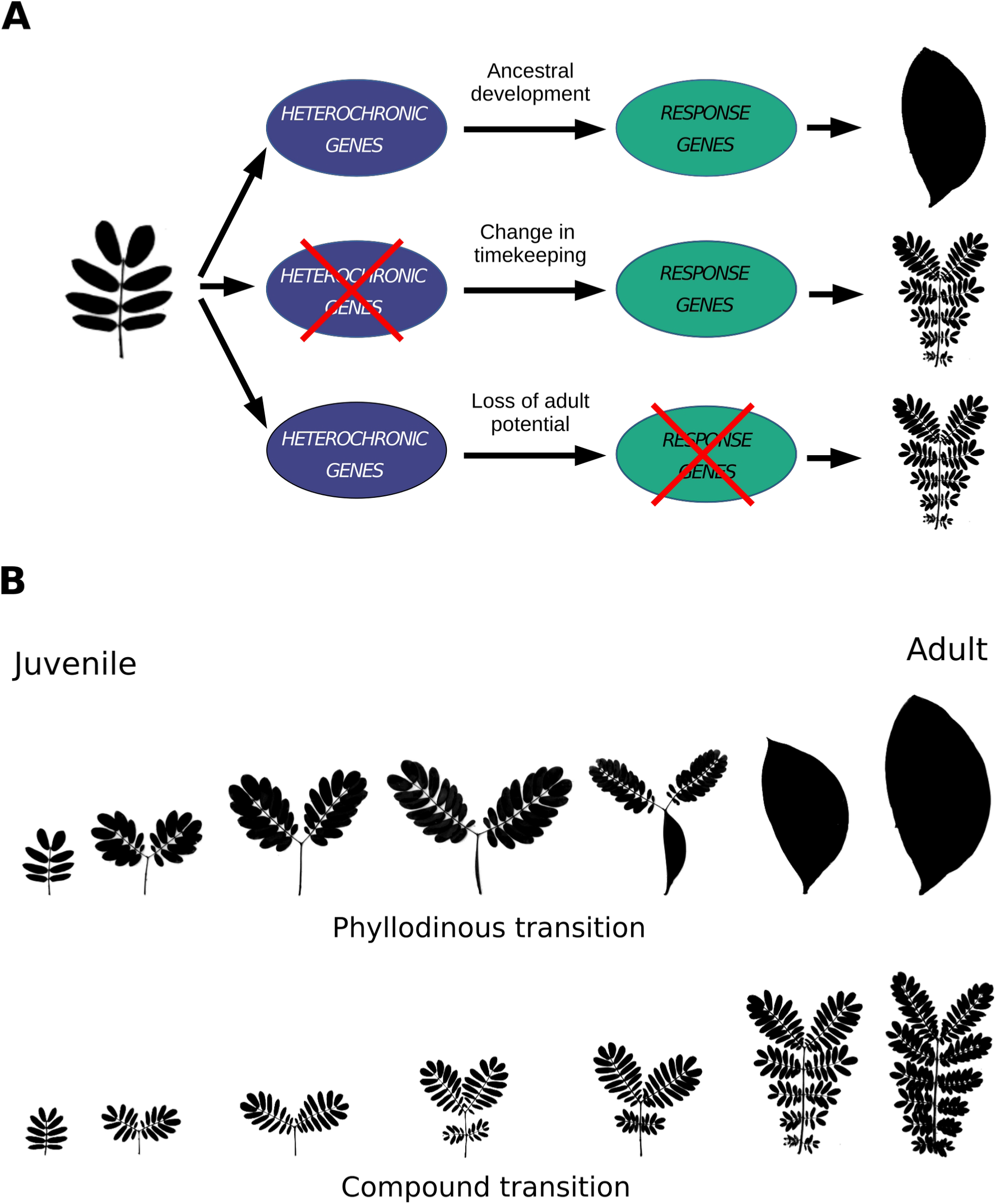
Potential genetic mechanisms for the evolution of neoteny. **(A)** In many plant and animal systems, genes that control the timing of developmental events are referred to as heterochronic. For timekeeping genes that promote juvenile development, loss of their expression would result in the retention of the juvenile phenotype (middle developmental progression). Conversely, loss of genes that respond downstream of the timekeeping pathways (or their ability to respond) would also result in neoteny (bottom developmental progression). We refer to the former mechanism as a “change in timekeeping” and the latter as a “loss of adult potential”. **(B)** Representative developmental transitions in leaf morphology found in the genus *Acacia*. Most *Acacia* species transition from a compound, juvenile morphology, to a simple, adult morphology (phyllodinous *Acacias*). Some species never produce the simple, adult leaf, and instead maintain the juvenile morphology (bipinnately compound *Acacias*).

Higher plants undergo two major developmental transitions during their post-embryonic development--the transition from juvenile to an adult phase of vegetative development (known as *vegetative phase change*) and the transition from a vegetative to a reproductive phase of development. Vegetative phase change is marked by changes in a variety of shared as well as species-specific traits, and can be quite subtle or very dramatic (Poethig and Fouracre, 2024). This transition is regulated by two closely related families of microRNAs, miR156 and miR157 (Wu and Poethig 2006; Rhoades et al. 2002; Poethig and Fouracre 2024). These miRNAs are expressed at high levels during embryo development and in the first few leaves produced after germination, and are then silenced. This decrease in miR156 and miR157 results in the derepression of their direct targets, a family of transcription factors known as the *SQUAMOSA PROMOTER PROTEIN BINDING-LIKE* genes (*SPL*) (Rhoades et al. 2002; Xu, Hu, Zhao, et al. 2016; He et al. 2018). *SPL* genes promote the transcription of genes that control the development of adult vegetative traits (Wang et al. 2011; Bergonzi et al. 2013; Zhou et al. 2013; Lawrence et al. 2021; Rubio-Somoza et al. 2014), and are thus the direct regulators of the adult vegetative phase.

Previous work found that a decline in the abundance of miR156 and miR157 is tightly correlated with the shift from juvenile to adult leaves in *Acacia* (Figure 1B), a large group of woody legumes found primarily in Australia and Southeast Asia (Wang et al. 2011). In *Acacia*, the first leaves produced by the shoot (hereafter “juvenile”, Figure 1B) are pinnately and bipinnately compound, which is the ancestral leaf type in legumes (D. Murphy et al. 2003; Renner et al. 2021). Most species in this genus then transition to producing a derived simple (i.e. undissected) leaf type called a phyllode (hereafter “adult”). The timing of this transition varies extensively within the genus, and is not necessarily correlated with a change in reproductive competence; for example, many *Acacia* species transition between alternative leaf morphologies within the first few weeks of development but do not flower for many years after this. Interestingly, some species of *Acacia* produce bipinnately compound leaves throughout their entire life cycle and flower in this condition (bipinnate species) (Brown et al., 2006; Miller et al, 2013). Traditionally, such species have been classified in two sections of *Acacia*, the Botrycephalae and the Pulchellae (Pedley 1986; Brown et al., 2006). Vegetative phase change in *Acacia* is therefore an excellent system for studying the molecular basis of heterochrony because this transition is obvious, its timing varies considerably within the genus, and it is likely controlled by known temporal (miR156 and miR157) and morphogenetic (*SPL*) genes.

Molecular phylogenies reveal that bipinnate species are nested within species that produce phyllodes (D. Murphy et al. 2003; D. J. Murphy 2008), and it has been hypothesized that they reflect a defect in developmental timing based on the observation that some members of this group, such as *A. latisepala*, sometimes produce phyllodes late in shoot development (Pedley 1986). However, there has been no comprehensive study of the phylogenetic relationship between species in which phyllode production is delayed and species that never produce phyllodes. This information is critical for determining whether the absence of phyllodes reflects a defect in developmental timing, or the inability to produce this derived leaf type.

To answer this question, we measured the timing of phyllode production in over 100 *Acacia* species and mapped this information onto a new and robust molecular phylogeny of the genus. We found that in many, but not all instances, bipinnate species are closely related to species that exhibit a delay in the bipinnate-to-phyllode transition. In three cases, we found that the prolonged production of bipinnate leaves is associated with either an increase or a delayed decline in the level of miR156, the master regulator of the juvenile-to-adult transition. We conclude that paeodomorphism in *Acacia* arises primarily from a change in developmental timing, potentially because of a change in the expression of miR156 or miR157.

## RESULTS

Although the transition from bipinnate leaves to phyllodes in *Acacia* was the basis for the conclusion that vegetative development in plants can be divided into juvenile and adult phases (Hildebrand 1875; Goebel 1889), there has been no systematic attempt to measure the timing of this transition in different *Acacia* species, or to map variation in timing of this transition onto the phylogeny of this genus. Therefore, to begin to study the mechanism of heterochrony in *Acacia*, we measured the time to produce the first phyllode (i.e the length of the juvenile phase) in 133 different accessions representing 106 phyllodinous species, grown in a single growth chamber. The average length of the juvenile phase in these species was 49.4 days, and ranged from 13 days in *Acacia confusa* to 195 days in *A. rubida* (Figure S1).

To determine how species with prolonged juvenile phases are related to bipinnate *Acacia* species, we generated double digest RAD-sequencing libraries (Peterson et al. 2012) for 201 individuals representing 148 named species (Table S1). Clustering across these individuals resulted in an average of 32,426 loci that passed coverage thresholds and a paralog filter. These loci represent an average of over 2.71 Mb per individual. We examined the phylogenetic support for the distribution of bipinnate clades by generating 10 supermatrices of concatenated loci using a different minimum number of individuals at a locus (“min”) and the maximum number of species with the same heterozygous position in a locus (“maxSH”), also known as the paralog filter (Table S2). In all trees, the bipinnate sections Botrycephalae and Pulchellae were recovered as distinct groups, near the crown and the base of the phylogeny, respectively (Figure 2 and Figure S2). Consistent with the results of Renner and colleagues (2021), we identified a third clade of bipinnate *Acacia* containing species such as *A. elata* and *A. terminalis* (Figure 2 and Figure S2). Additionally, there were other bipinnate species grouped in clades not previously supported (*A. gilbertii*) or examined by other molecular phylogenies (*A. chrysotricha*).

**Figure 2.**
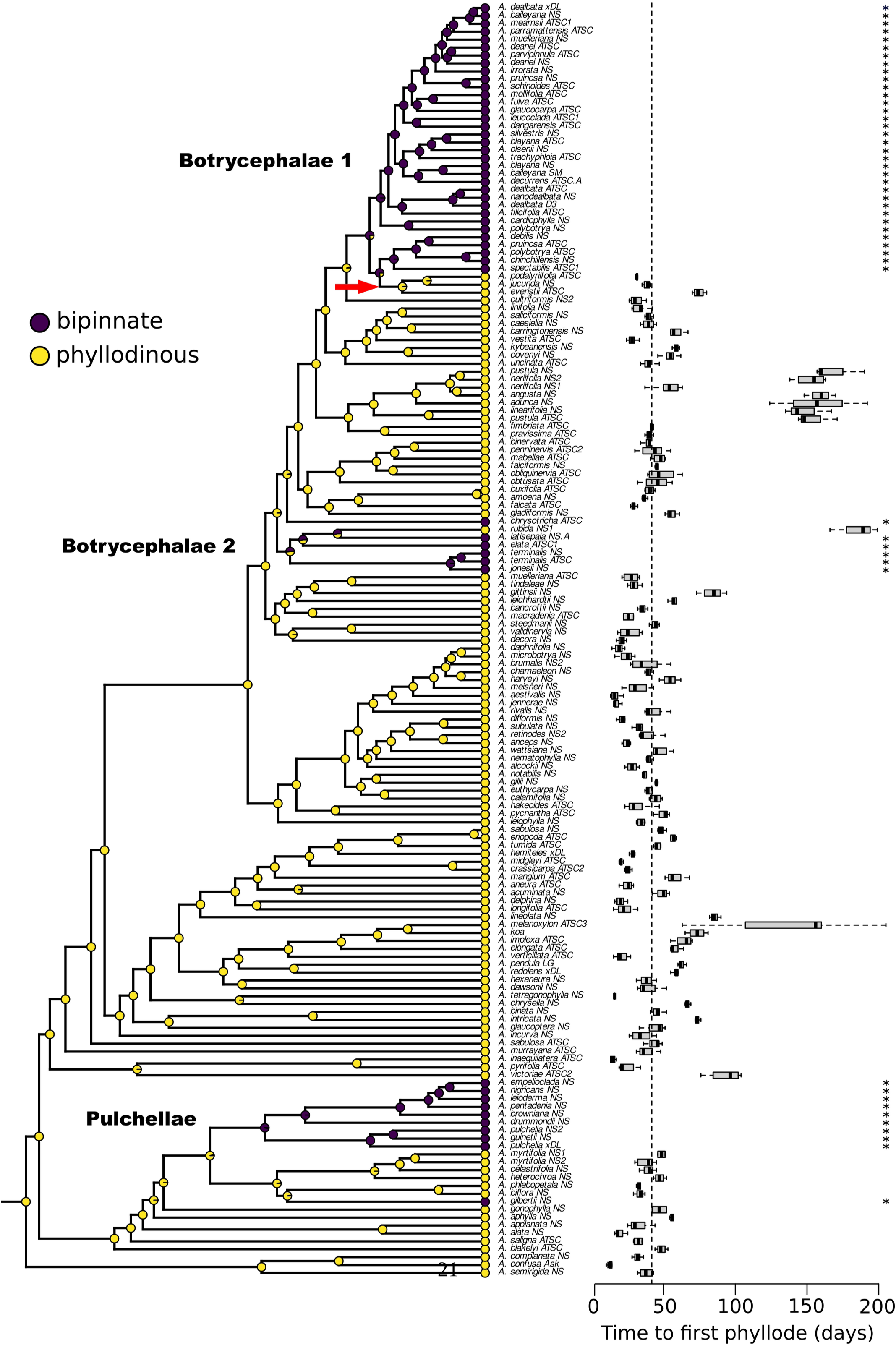
Evolution of neoteny in *Acacia*. The phylogeny represents an ancestral character state reconstruction of neoteny (retention of the juvenile morphology into adulthood, i.e. bipinnately compound leaves). Proportion of a circle’s color at internal nodes represent the posterior probabilities of each state using stochastic character mapping. The bold labels near internal branches of the tree denote historic classifications based on morphology of bipinnate groups. The notation of groups as either “Botrycephalae 1” or “Botrycephalae 2” is to highlight their polyphyly. The red arrow highlights a predicted reversion to the adult morphology. The boxplot to the right of the tree plots the median time to produce the first phyllode in phyllodinous species grown under common conditions. The dashed vertical line represents the median time to produce phyllodes in all phyllodinous accessions tested. The “*” represents the fact that bipinnately compound species by definition do not produce phyllodes.

To evaluate how loss of phyllodes evolved within these groups, we trimmed our data matrix to include only one individual per species (Figure S8) and generated a chronogram for the genus. Ancestral character state mapping of the phyllodinous and bipinnate states onto this tree revealed an estimated 7 transitions to the bipinnate state, and 2 transitions to the phyllodinous state (Figure 2 and Figure S9). For section Pulchellae, we identified two transitions to the bipinnate state. One of these transitions is represented by multiple species, including *A. pulchella*, and the other is represented by the single species, A*. gilbertii*. This grouping was consistent and well supported across all phylogenies (Figure S3 & S4). There was no evidence of reversion to the phyllodinous state within this section.

We found two distinct bipinnate clades in the Botrycephalae (Figure 2, Botrycephalae 1 and 2). The smaller of the two clades contained *A. elata*, *A. jonesii*, *A. terminalis*, *A. latisepala*, and *A. rubida*. *A. rubida* is phyllodinous (Figure 2, Botrycephalae 2), and our character state reconstruction supports the conclusion that this species represents a re-acquisition of the phyllodinous state. In all trees examined, the bipinnate *A. latisepala* and phyllodinous *A. rubida* were sister species with high bootstrap support (Figure S3 & S5). Additionally, the monophyly of this clade was supported by trees generated from all supermatrices, except for the 150-3 matrix, which had the fewest number of loci. The other bipinnate clade consists of 37 species, including such species as *A. spectabilis* and *A. mearnsii* (Figure 2, Botrycephalae 1). Consistent with other studies (Renner et al. 2021), we found a small clade of phyllodinous species (*A. podalyriifolia*, *A. jucunda*, and *A. everistii*) nested within this group (Figure 2, red arrow). Our character reconstruction suggests that this group is a reversion to the phyllodinous state from a bipinnate ancestor (Figure 2). This grouping was also well supported by all of our phylogenies (Figure S3 & S7).

If the retention of juvenile leaf morphology is the result of changes in timekeeping genes rather than the loss of the ability to produce adult traits (Figure 1A), then species that have abnormally long juvenile phases are expected to be phylogenetically related to permanently juvenile species. As mentioned above, a good example of this is *A. rubida,* which was the slowest switching phyllodinous species we tested, and is closely allied with bipinnate species. This observation suggests that these bipinnate species acquire this state by more severe delays in developmental timing. Phylogenetic evidence for reversion to a phyllodinous phenotype also suggests that the bipinnate state is consistent with a change in developmental timing. The large bipinnate clade containing the phyllodinous species *A. podalyriifolia*, *A. jucunda*, and *A. everistii* is an example of this (Figure 2, Botrycephalae 1).

To test the prediction that bipinnate species are indeed juvenilized species, we measured the abundance of miR156 and miR157 in species with different temporal patterns of vegetative phase change. We first examined intraspecific differences in the timing of phyllode production in two accessions of *A. melanoxylon*. One of these accessions began producing adult leaves, on average, 65 days after planting (Figure 3A, “fast”) whereas the other accession required more than 120 days of growth (Figure 3A, “slow”). Consistent with the results for *A. confusa* and *A. colei* (Wang et al. 2011), miR156 and miR157 declined in association with the switch between bipinnate leaves and phyllodes in both accessions. However, both miR156 and miR157 declined more slowly in the “slow” switching accession than in the "fast" switching accession.

**Figure 3.**
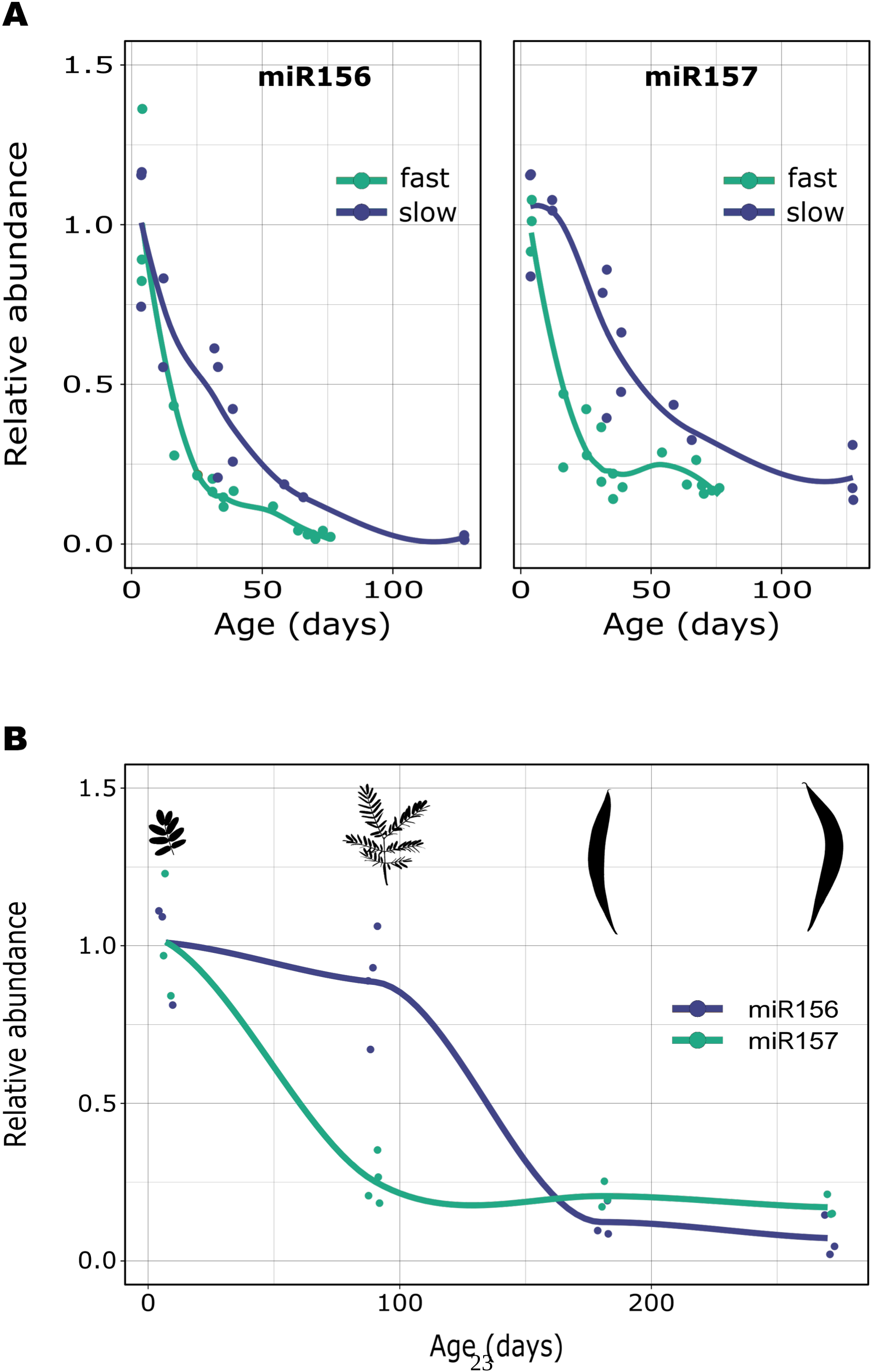
Delays in the timing of adult leaf development are associated with difference in miR156 and miR157 abundance. (**A**) miR156 and miR157 levels in two accessions of *A. melanoxylon*. The “fast” accession switches from juvenile to adult leaves within 3 months whereas the “slow” accession requires more than 5 months. (**B**) miR156 and miR157 levels in *A. rubida*, a species that transitions from juvenile to adult leaves after 6-8 months of growth. The leaf silhouettes are representative of the average leaf morphology of the most recently produced leaves at each time point.

We next examined if these microRNAs are associated with interspecific variation in the timing of the bipinnate-to-phyllode transition. We first measured the abundance of miR156 and miR157 in 3 bipinnate members of the Botrycephalae and 3 related phyllodinous species over the course of 3 months (a period that exceeds the average time to produce phyllodes for the entire genus) (Figure 4). In all six species, miR157 declined to similar levels over this time period. However the level of miR156 in bipinnate species was significantly higher at three months than in phyllodinous species (Figure 4B). Together, these results suggest that at least two instances of paedomorphism in the genus *Acacia* have evolved by changes in developmental timing, and this delay in development is mediated by changes in genes that regulate the timing of the juvenile to adult transition in leaf morphology in plants.

**Figure 4.**
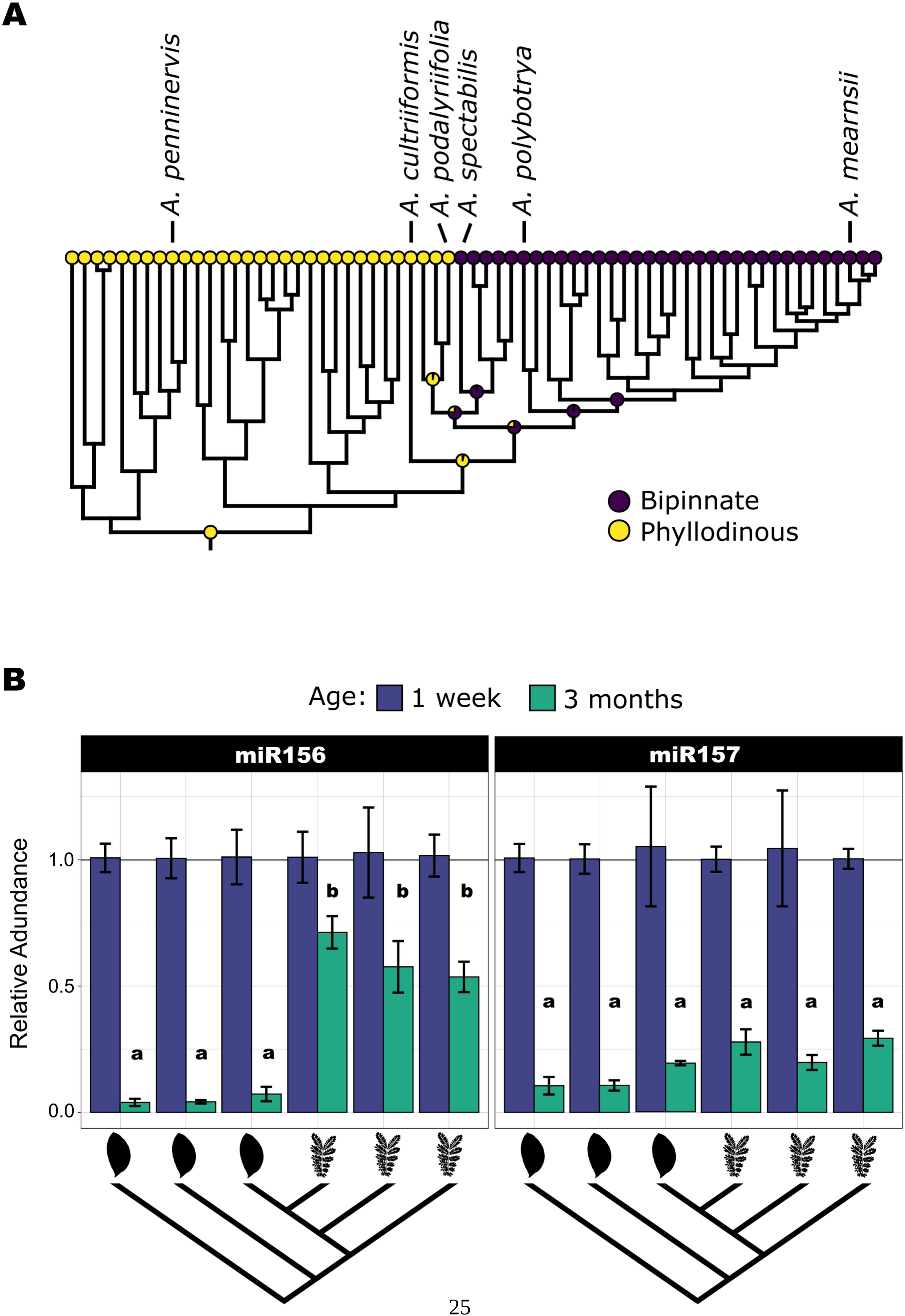
Paedomorphic leaf morphology in the subgenus Botrycephalae is associated with a difference in miR156 abundance. (**A**) Phylogenetic position of species examined in B, and their associated adult leaf morphology (yellow = phyllodes; purple = compound leaves). Ancestral state reconstructions are from Figure 2. (**B**) Levels of miR156 and miR157 in the first leaves at 1 week post-germination, and in the most recently fully expanded leaves at 3 months post-germination. The phylogeny at the bottom of the panel is a simplified representation from panel A showing the morphology of the adult leaves (either phyllode or bipinnate). Data are means of 3-6 biological replicates ± SEM. Letters above the 3-month bars indicate a significant difference at *P*<0.05, Tukey’s HSD test. Groups with different letters are significantly different. Species from left to right are, *A. penninervis*, *A. cultriformis*, *A. podalyriifolia*, *A. spectabilis*, *A. polybotrya*, *A. mearnsii*.

As discussed above, *A. rubida* is a slow switching species that is phylogenetically close to constitutively juvenile species such as *A. elata* and *A. latisepala* (Figure 2, “Botrycephalae 2”). Furthermore, ancestral character state reconstruction suggests it is a reversion to phyllode production (Figure 2). In contrast to our findings for *A. melanoxylon* accessions, and previous reports for *A. confusa* and *A. colei* (Wang et al. 2011), we found that the decline in miR156 and miR157 are decoupled, with miR157 declining within the first 3 months of growth (Figure 3B) similar to the patterns observed in phyllodinous and bipinnate species (Figure 4B). Comparatively, the decline in miR156 was delayed to over 6 months of growth, consistent with the relatively late production of phyllodes (Figure 3B) and a pattern similar to miR156 in bipinnate species belonging to Botrycephalae 1 (Figure 4B). These findings suggest that a delay in miR156 may be responsible for the delay in adult leaf production in *A. rubida*, and may also explain the bipinnate character of sister species such as *A. latisepala*. Unfortunately, we were unable to grow enough plants from bipinnate species in this clade to directly test this prediction.

## DISCUSSION

While heterochronic shifts in major life history transitions are common and often happen repeatedly in many lineages, the mechanism of these morphological shifts remains unclear. Indeed, this phenomenon may not actually reflect a change in developmental timing (Cubo et al. 2000); retention of juvenile features into adulthood could simply result from the loss of genetic pathways that are responsible for the production of adult traits. For example, in plants, wood formation is absent during early seedling development, and is generated by secondary growth of the vascular cambium later in shoot development (Onyenedum and Pace 2021). In monocots, this adult phase of vascular development is absent, and plants retain the juvenile phase of vascular development throughout their entire life cycle. Interestingly, monocots lack genes necessary for secondary xylem formation (Roodt et al. 2019), suggesting that they are paedomorphic by loss of genes necessary for adult trait development (Onyenedum and Pace 2021). Alternatively, neoteny could result from changes in the genetic pathways that control developmental timing. In this case, neotenous species would retain the ability to produce adult traits, but be prevented from expressing this potential. Phylogenetic reversion to the ancestral adult phase of development would not necessitate the re-acquisition of adult developmental pathways, but simply a change in developmental timekeeping genes.

Our results suggest that changes in developmental timing have played a major role in species differences in *Acacia*. This conclusion is supported by the close phylogenetic relationship between species with delayed phase change and species that never undergo this developmental transition, and by the observation that miR15—6--a miRNA that promotes the juvenile phase—is expressed at elevated levels (i.e. juvenile-like levels) in plants in which vegetative phase change is either delayed, or does not occur. Species that retain juvenile traits throughout their development have also been identified in *Eucalyptus* (Boland et al, 2006), several species in the Cupressineae (Schaffalitzky de Muckadell, 1954) and many species native to New Zealand (Godley, 1985). In *Eucalyptus globulus*, variation in the timing of.vegetative phase change maps to a QTL that includes *EglMIR156.5*, and this gene is expressed at lower levels in F2 progeny with precocious phase change than in siblings with normal phase change (Hudson et al. 2014). It remains to be determined if inter-specific variation in the timing of vegetative phase change in *Eucalyptus* or other genera is associated with variation in the expression of miR156 or miR157, but it would not be surprising if this were true. In any case, the existence of multiple cases of paedomorphism within these and other plant genera, along with a potential molecular mechanism for this phenomenon, make these ideal systems for research on the mechanism of heterochrony in organismal evolution.

In *Arabidopsis*, the decrease in the abundance of miR156 during vegetative phase change is mediated by the acquisition of the repressive epigenetic mark, H3K27me3, at two genes that produce the majority of miR156 in this species (Xu, Hu, Smith, et al. 2016). How timing of this process is determined endogenously (i.e. by the propensity of a miR156/miR157 locus to acquire silencing marks), and how it is modified by environmental conditions that increase or decrease the duration of the juvenile phase remains an open question (Cheng et al. 2021; Gao et al. 2022; Poethig and Fouracre 2024). This process depends on the activity of the factors that mediate the deposition of H3K27me3 (and other epigenetic marks), and on the sensitivity of miR156/miR157 genes to these factors. Natural variation in the expression of these genes can therefore be the result of either a change in their cis-regulatory sequences, or a change in the factors that bind to these sequences. These scenarios make different predictions about the genetic basis of variation in the level of miR156/miR157. Changes in cis-regulatory sequences are expected to affect the expression of a single gene, whereas changes in trans-acting regulatory factors are expected to affect the expression of all the genes regulated by these factors--e.g. all the miR156/miR157 genes that are temporally regulated by these factors. For example, intra-specific variation in the timing of vegetative phase change in *Eucalytus globulus* is likely attributable to a cis-regulatory change in *EglMIR156.5,* because sequencing of the stem-loop region did not reveal polymorphisms between normal and precocious individuals or expression differences at other *MIR156* genes (Hudson et al. 2014).

We were unable to examine the expression of miR156/157 in all of the *Acacia* species representing independent evolution of the paedomorphic phenotype due to practical issues related to seed availability and germination issues (e.g. section Pulchellae). However, the lack of evidence for an association of bipinnate species in section Pulchellae with delays in developmental timing in allied species, as well as the lack of evidence for reversion to phyllode production in this section, suggests that this phenotype may reflect a loss of the ability to produce phyllodes. In this regard, it is significant that other vegetative traits are regulated in an age-dependent manner in the Pulchellae, including the length of the petiole and the appearance of axillary spines during seedling development (Leichtyunpublished observation). Future analyses of the expression of miR156/miR157 and their *SPL* targets may help to resolve this question.

When vegetative phase change is marked by changes in multiple morphological traits, different traits can appear at slightly different times and can become decoupled, producing plants that are a mixture of juvenile and adult traits (Hackett and Murray, 1992; He et al. 2018; Leichty and Poethig 2019; Lawrence et al. 2021). This suggests that each trait has a unique threshold or sensitivity to levels of miR156/miR157. This variation could be mediated by differences in transcriptional activity of *MIR156*/*MIR157* genes in different organs, or by the varying sensitivity of different *SPL* genes to these miRNAs. Changes in these response pathways are likely to be a large source of heterochronic variation, but in plants these pathways are more difficult to characterize than measuring levels of well defined upstream regulators of developmental phase transitions (e.g. miR156 or *FLC*) (Cartolano et al. 2015).

The most detailed studies of neoteny in animals have been conducted in amphibians, where thyroid hormone (TH) mediates the process of metamorphosis, the transition between juvenile and adult phases of somatic development (Shi 2000). The salamander, *Ambystoma mexicanum* never undergoes metamorphosis and become reproductively mature in a somatically juvenile state. QTLs associated with neoteny in this species display reduced TH sensitivity (Voss et al. 2012). Similarly, in *Eurycea tynerensis,* thyroid hormone receptors (TRs) from neotenous populations have reduced TH sensitivity when compared with metamorphic relatives (Aran et al. 2014). In other instances, such as neotenous *Necturus* salamanders, TRs have been shown to be functional and expressed in target tissues, but some of their downstream target genes appear to have lost responsiveness (Safi et al. 2006). Together, these findings suggest that the heterochronic shifts in neotenous salamanders often evolve by modifications of TH response and possibly to changes in TH abundance (similar to routes 2 and 3 in Figure 1A), although the molecular basis of these phenotypes has not been established.

Our results suggest that neoteny has also played a major role in plant evolution, and provide a plausible molecular mechanism for this phenomenon in *Acacia*. The availability of whole genome sequences for several *Acacia* species, as well as a protocol for the genetic transformation of at least one of these species (McLay et al. 2022; Massaro et al. 2024), opens the way for further molecular genetic investigations of this important mechanism of organismal evolution.

## METHODS

### Plant material and growth conditions

All plants were germinated in petri dishes on filter paper after clipping the seed coat. Seeds were then transferred to 3” or 1 gallon pots containing Fafard-2 potting mix, depending on the experiment and duration of growth. For the analysis of phyllode timing, plants were grown in a Conviron chamber maintained at 24°C, with 16 hrs. light/8 hrs. dark, and 190-220 μmol m^-2^ s^-1^ using white fluorescent lights. Plant age was calculated from date of germination. Plants were periodically inoculated with *Amblyseius cucumeris* to prevent thrip infestation. Plants sampled for smRNA abundance were grown in a greenhouse with supplemental light. Leaves were collected at full expansion, from 5-7 hours after sunrise.

### Preparation of RAD-seq libraries and sequencing

Double digest restriction associated DNA sequencing (ddRAD-seq) libraries were prepared according to the original method (Peterson et al. 2012). Briefly, genomic DNA was extracted using the DNeasy Plant Kit (Qiagen) and DNA was eluted into EB buffer (Qiagen). 500 ng of DNA was double digested with HindIII-HF and MfeI-HF (NEB) for 4 hours at 37°C at a volume of 50 ul. Digests were confirmed on 0.5% agarose gels. Half of the digest was used for adapter ligation with 6.5 pmol of each annealed adapter in a total volume of 50 ul. At this stage, 12 samples with unique P1 barcodes were pooled and cleaned using a 1X concentration of AMPure XP beads (Beckman Coulter). One microgram of the resulting pools was size selected on a Pippen Prep (Sage Science) for an insert size of 350 bp +/-12% using 1.5% gels. Size selected pools were then amplified and indexed in 5-10 replicate PCR reactions using Q5 polymerase (NEB). PCR reactions were then combined and bead-cleaned as above, and eluted into a final volume of 30 ul. Final concentrations were determined using a Qubit fluorometer, and sequencing-pools of 3-5 cleaned PCR pools were combined at equal mass (a total of 36-60 samples). Before Illumina sequencing, libraries were quality checked by blunt end cloning and Sanger sequencing using the CloneJET PCR Cloning Kit (Thermo Scientific). Libraries were sequenced at the University of Pennsylvania Next Generation Sequencing Core (Philadelphia, PA) on a HiSeq 2500 with 100SE format.

### Phylogenetic analyses

Pyrad (version 3.0.66) was used for identifying nucleotide variants in the ddRAD-seq data (Eaton 2014). Reads were demultiplexed and filtered using default settings. Clustering was performed at 85% similarity and a minimum coverage of 6 for any given locus. A variety of supermatrices were evaluated by changing the maximum number of individuals with a shared heterozygous position (i.e. the paralog filter) and the minimum number of samples for a cluster (Table S2).

For each supermatrix, maximum likelihood trees were generated using RAxML v8.2 (Stamatakis 2014) with the GTRGAMMA model of rate heterogeneity using the rapid bootstrapping mode with 100 searches. *Paraserianthes lophanta* was used as an outgroup. For the final tree used for character mapping, we removed multiple accession representing a single species. Specifically, we eliminated individual libraries corresponding to the same species when they grouped together on the min90_maxSH3 tree, leaving only 1 representative per species. In cases where individuals labeled as the same species grouped with another species they were left in the tree as long as the total distance between them was greater than 0.0015 (this was the average distance between intraspecific pairs across the entire phylogeny). A full list of samples used in this study can be found in Supplemental Table S1. The final chronogram was estimated from the resulting tree (162 tips) using a Penalized Likelihood Approach with the *chronos* function in APE (Paradis, Claude, and Strimmer 2004) using R Statistical Software (v4.4.1; (R Core Team 2024)). The final lambda value of 0.1 and “correlated” model were selected based on log-likelihood values. The root age was set to 23 Ma following previous work (Macphail and Hill 2001; Miller et al. 2013; Renner et al. 2020).

Ancestral character reconstruction was implemented using phytools v1.9.1 (Revell 2012; 2023). For discrete character analysis Maximum Likelihood was used for fitting ER, ARD, and irreversible models, and AIC was used for model selection. Stochastic character mapping was used with the best rates model using 1000 trees.

### qPCR of smRNAs

Total RNA was extracted using the Spectrum^TM^ Plant Total RNA Kit (Sigma-Aldrich). cDNA was synthesized using Invitrogen SuperScript III following the manufactures specifications and previously published methods and primers for stemloop reverse transcription (Table S3) (Varkonyi-Gasic et al. 2007; Leichty and Poethig 2019). Platinum Taq (Invitrogen) was used with the Roche Universal Hydrolysis Probe #21, and a three step amplification protocol with previously published primers (Leichty and Poethig 2019). Relative measures of abundance were calculated using the 2^-ΔΔCt^ method (Livak and Schmittgen 2001) using miR159 and miR168 as endogenous controls (Leichty and Poethig 2019). R Statistical Software (v4.4.1; (R Core Team 2024)) and ggplot2 was used for plotting qPCR data with the geom_smooth() function for all curves (Wickham 2016) and linear models and tests were fitted using the base functions lm(), aov(), and TukeyHSD().

## DATA AVAILABILITY

Raw sequencing reads have been submitted to Genbank under the bioproject: PRJNA988539. Phenotypic data and tree files are available on DRYAD.

## ACKNOWLEDGMENTS

We thank the University of Pennsylvania Next-Generation Sequencing Core for help with sequencing and the University of Arizona Desert Legume Program for donation of some seed accessions.

## Supplemental Figures

**Figure S1.**
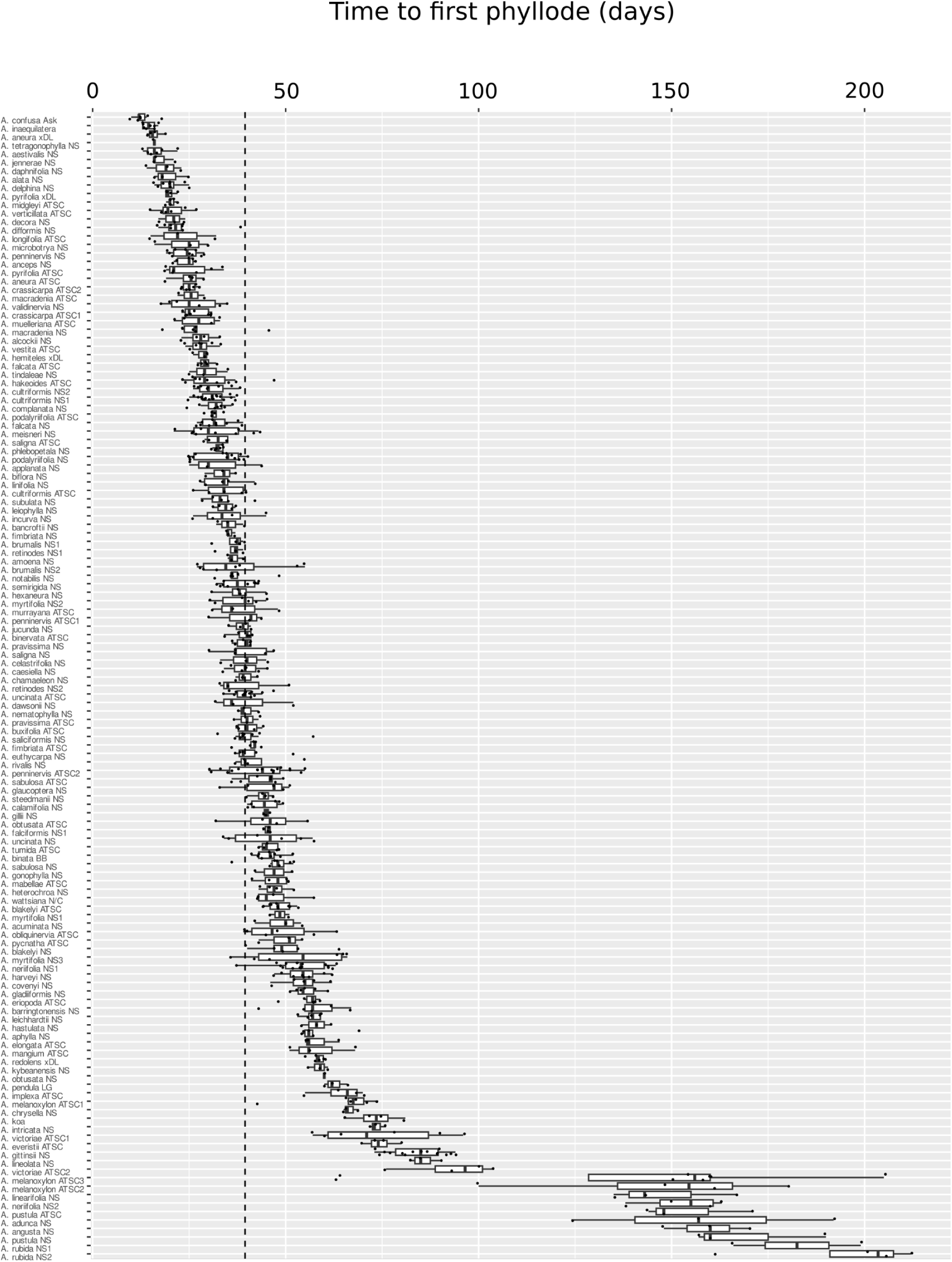
Length of the juvenile phase in phyllodinous *Acacia* species and accessions. The vertical line represents the median length across all measured species.

**Figure S2.**
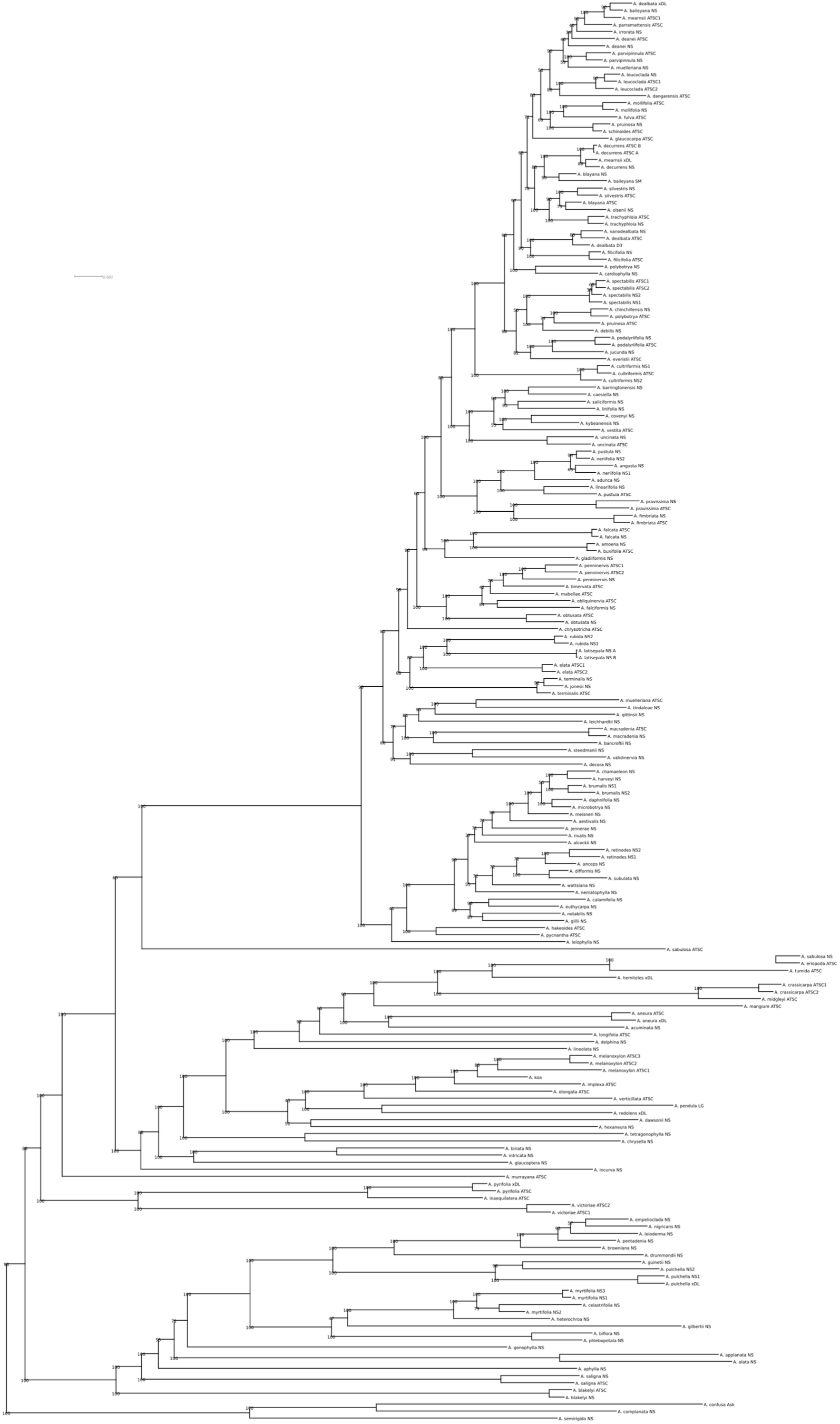
Full Maximum Likelihood phylogeny of all 201 libraries generated from the min90-maxSH3 supermatrix. Branch labels represent bootstrap support. *P. lophant*a has been removed from the tree.

**Figure S3.**
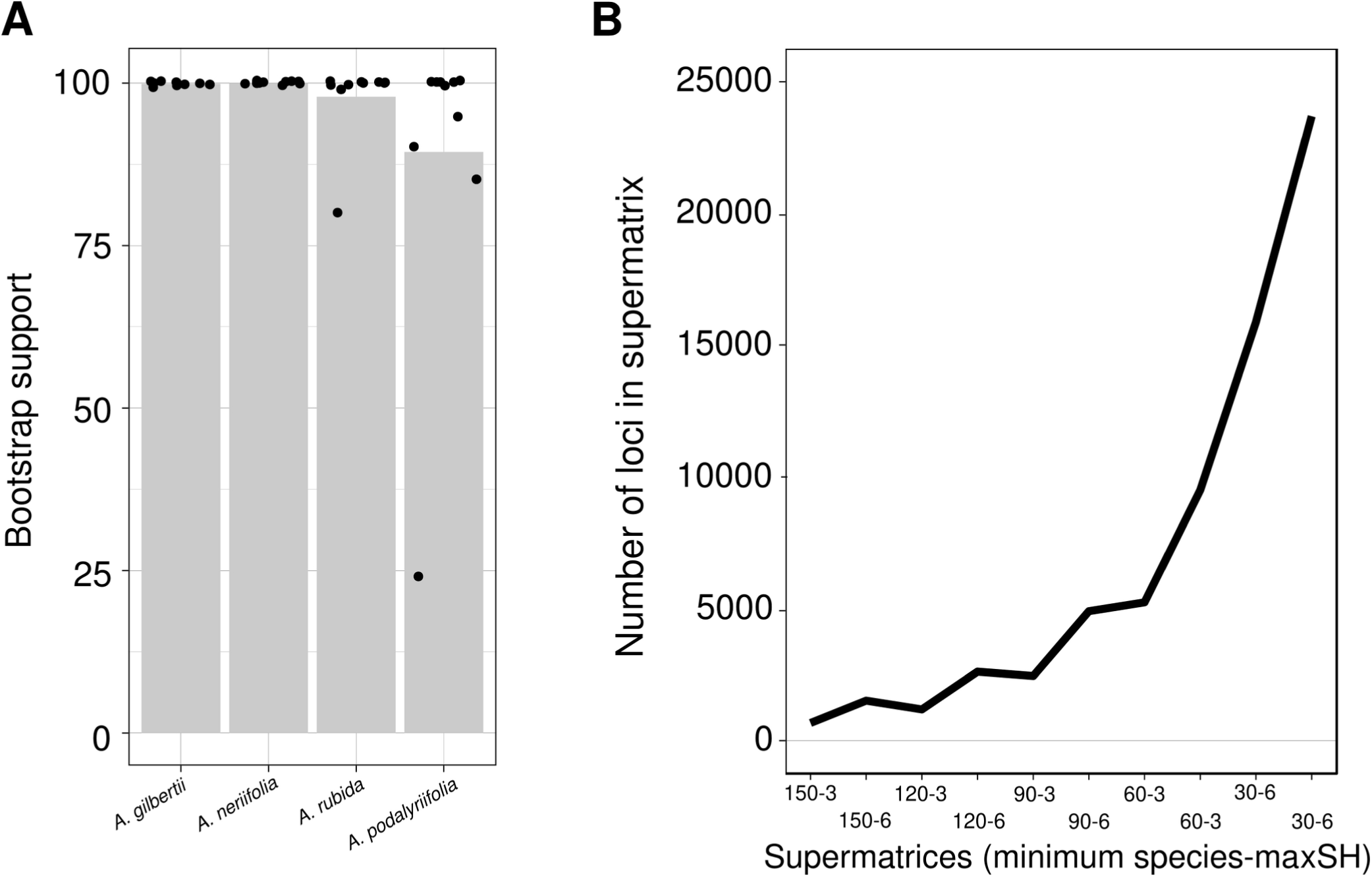
Assessment of phylogenetic uncertainty for key clades in the phylogenetic datasets. **(A)** Bootstrap support for four different clades: *A. gilbertii* grouped with phyllodinous species (including *A. myrtifolia*), a grouping of *A. neriifolia* with other slow switching phyllodinous species, *A. rubida* with *A. latisepala*, and a group of phyllodinous species (including *A. podalyriifolia*) as sister to a group of non-phyllodinous species (including *A. spectabilis* and *A. debilis*). See Figures S4-S7 for phylogenetic placements. **(B)** The number of loci in each supermatrix as a function of decreasing minimum species and paralog filter thresholds.

**Figure S4.**
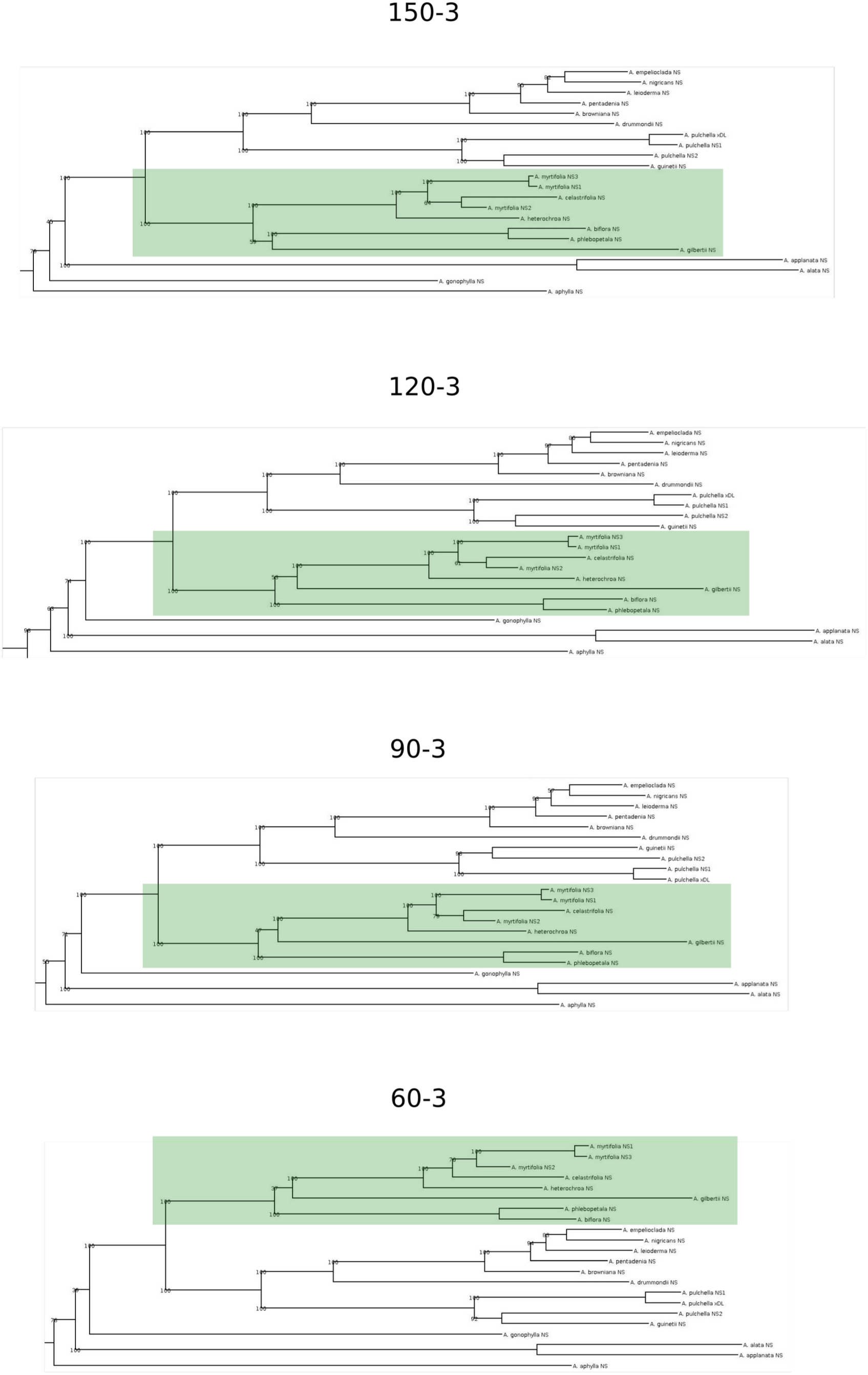
Phylogenetic position of *A. gilbertii*, a non-phyllodinous species nested within phyllodinous species. Trees generated from all supermatrices always placed *A. gilbertii* within this group, and bootstrap support was always 100.

**Figure S5.**
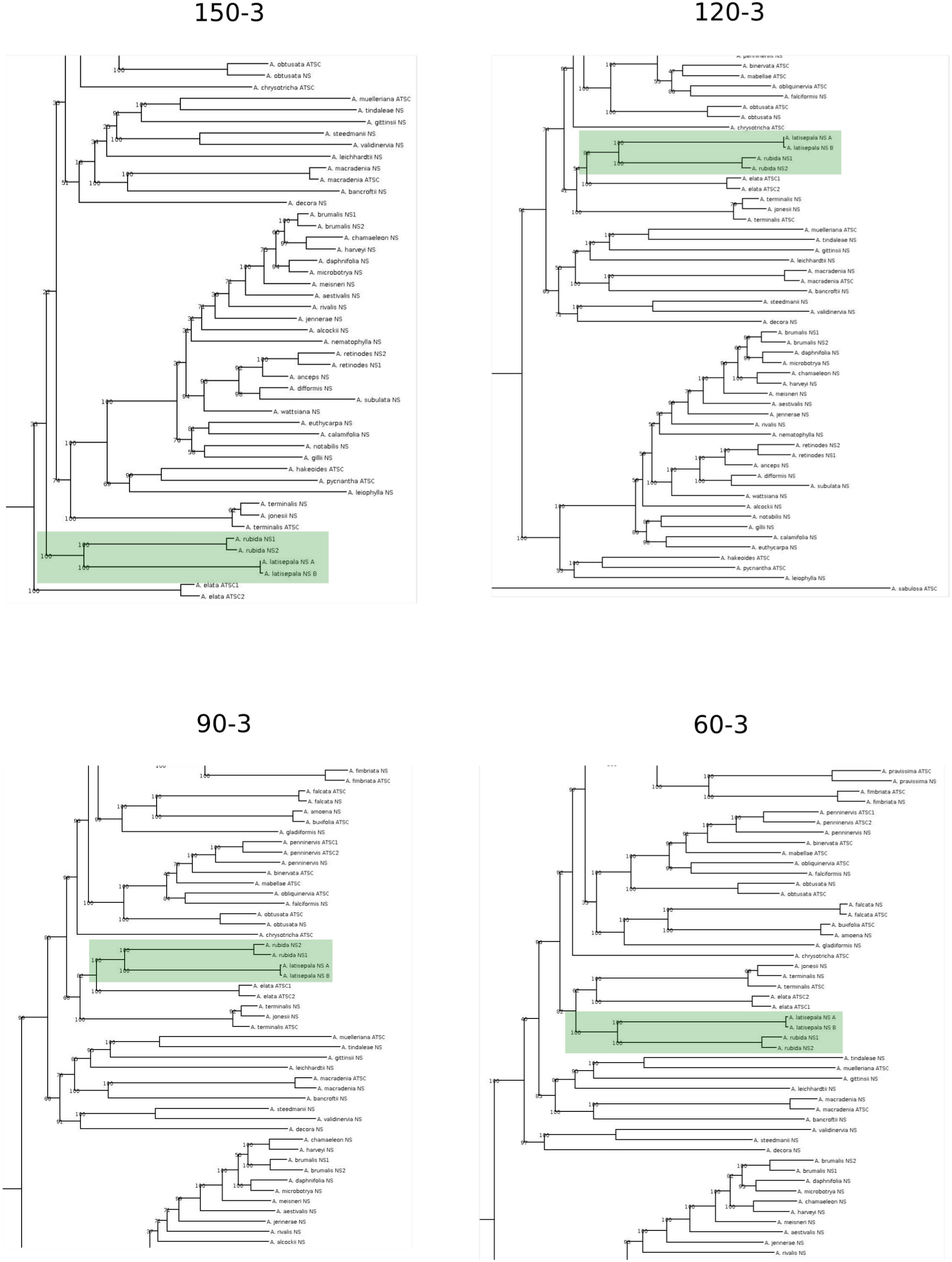
Phylogenetic position of *A. rubida* relative to *A. latisepala*. Best trees generated from all supermatrices always placed *A. rubida* and *A. latisepala* as sister species. Bootstrap support ranged from 80-100. *A. rubida* is a slow switching phyllodinous species, while our accessions of *A. latisepal*a never transitioned and were classified as non-phyllodinous. These two species where nearly always clustered with other non-phyllodinous species such as *A. elata* and *A. terminalis*. The one exception being in supermatrices with small numbers of loci (e.g. the 150-3 supermatrix) where these species where separated into early diverging lineages relative to their positions in trees with larger numbers of loci (e.g. 120-3, 90-3, 60-3, 30-3).

**Figure S6.**
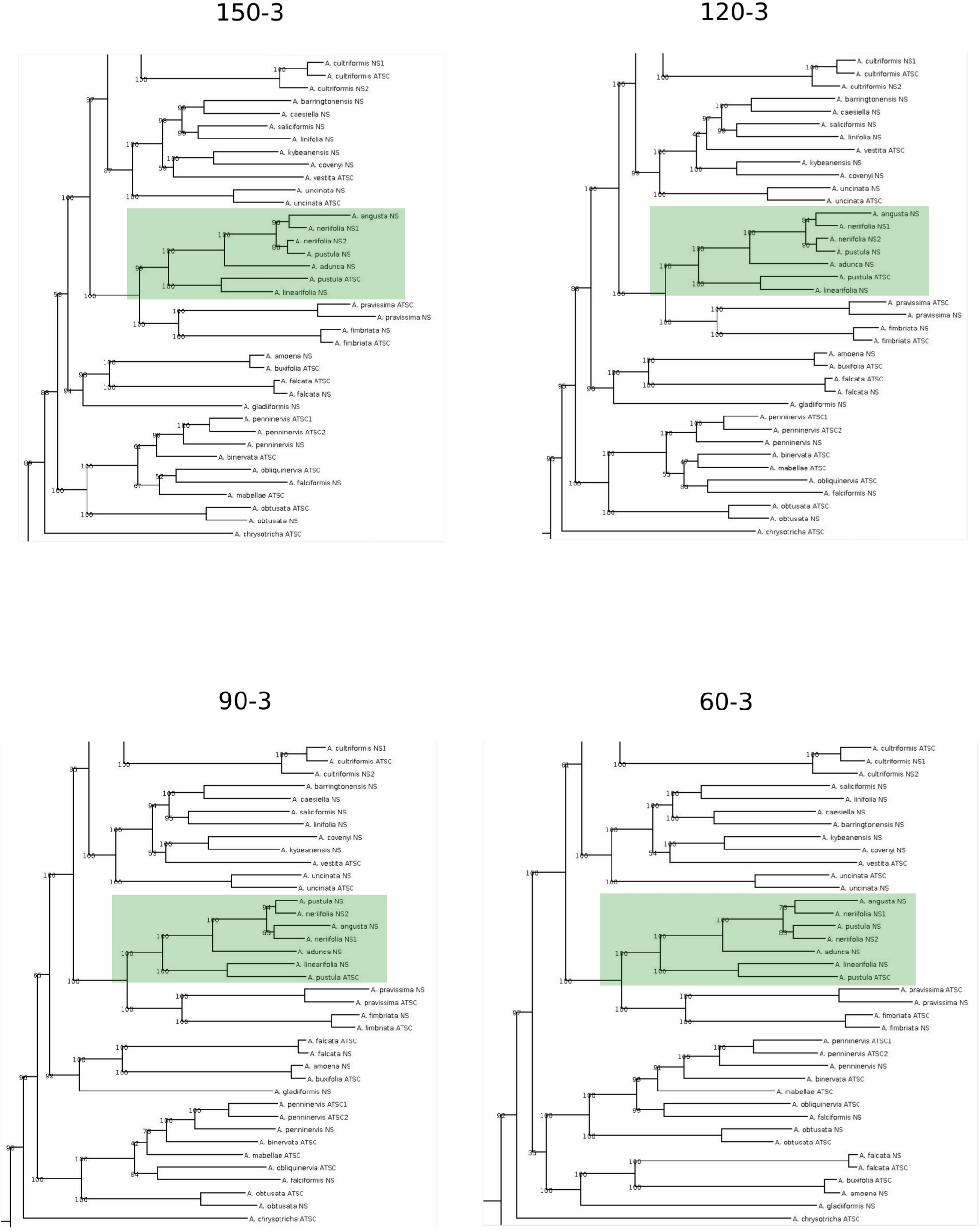
Phylogenetic position of a group of slow switching phyllodinous species that includes *A. neriifolia*. Trees generated from all supermatrices always placed these species together, with this grouping having bootstrap support from 99-100.

**Figure S7.**
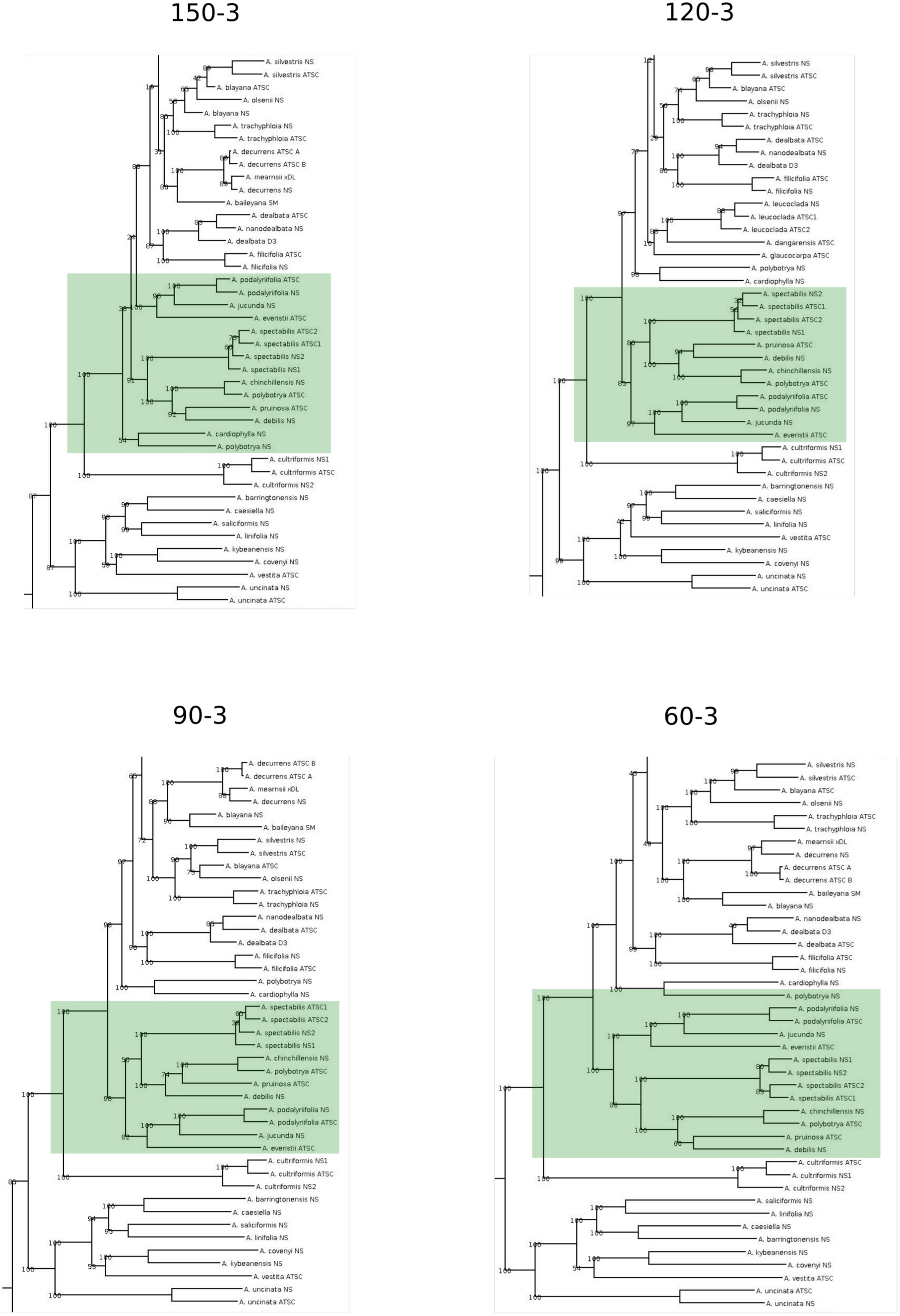
Phylogenetic position of phyllodinous species *A. podalyriifolia*, *A. jucunda*, and *A. everistii* relative to the core Botrycephalae. Trees generated from all supermatrices always placed this *A. podalyriifolia* clade within a clade of non-phyllodinous Acacias with this grouping always having a bootstrap support of 100. Most trees placed this clade as sister to a group containing *A. spectabilis* and *A. debilis*, with bootstrap support ranging from 24-100.

**Figure S8.**
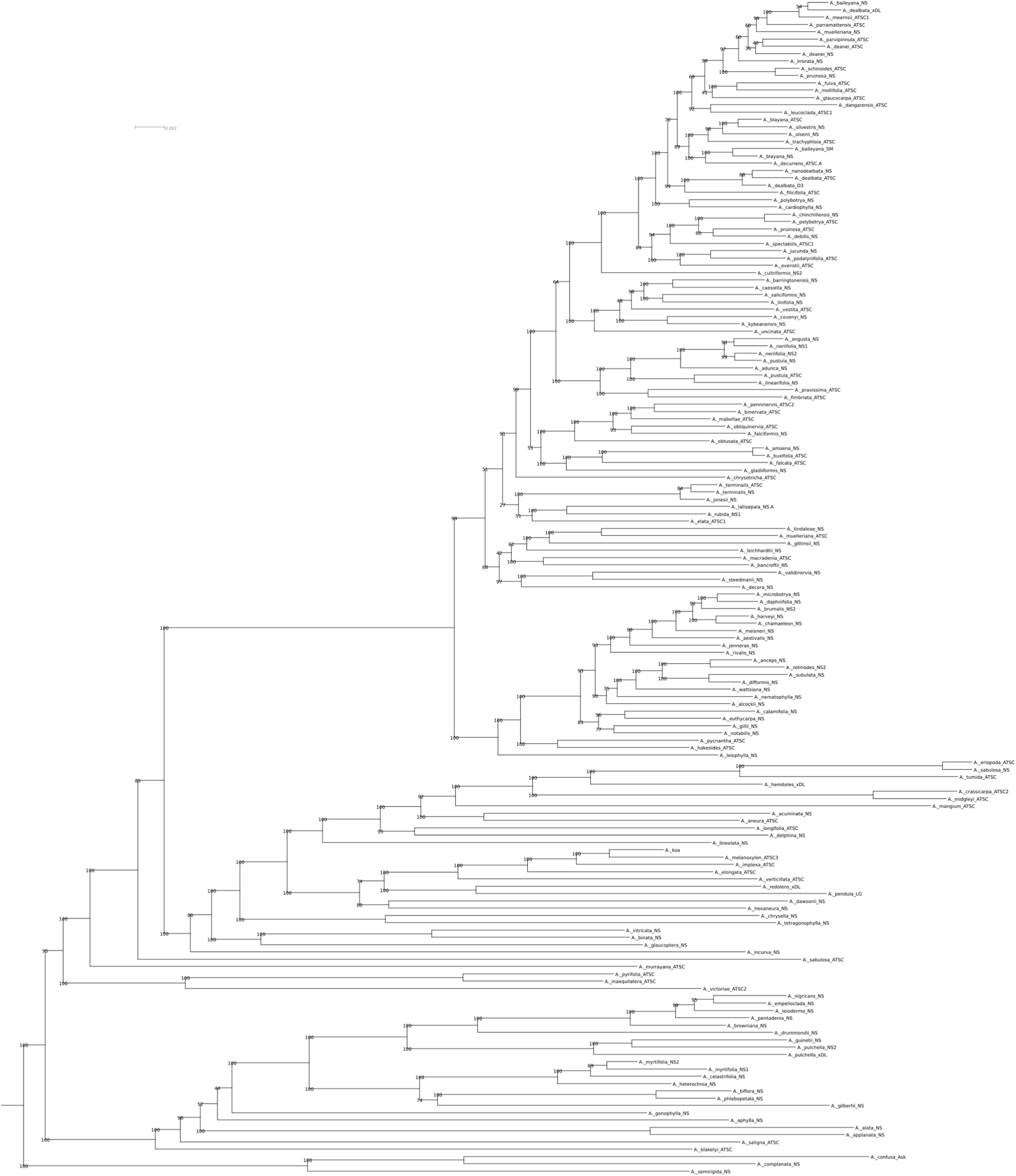
Phylogeny used for ancestor character reconstruction (Figure 2). Multiple individuals for a single species were removed from the full phylogeny (see methods for description). *P. lophanta* has been trimmed from this tree.

**Figure S9.**
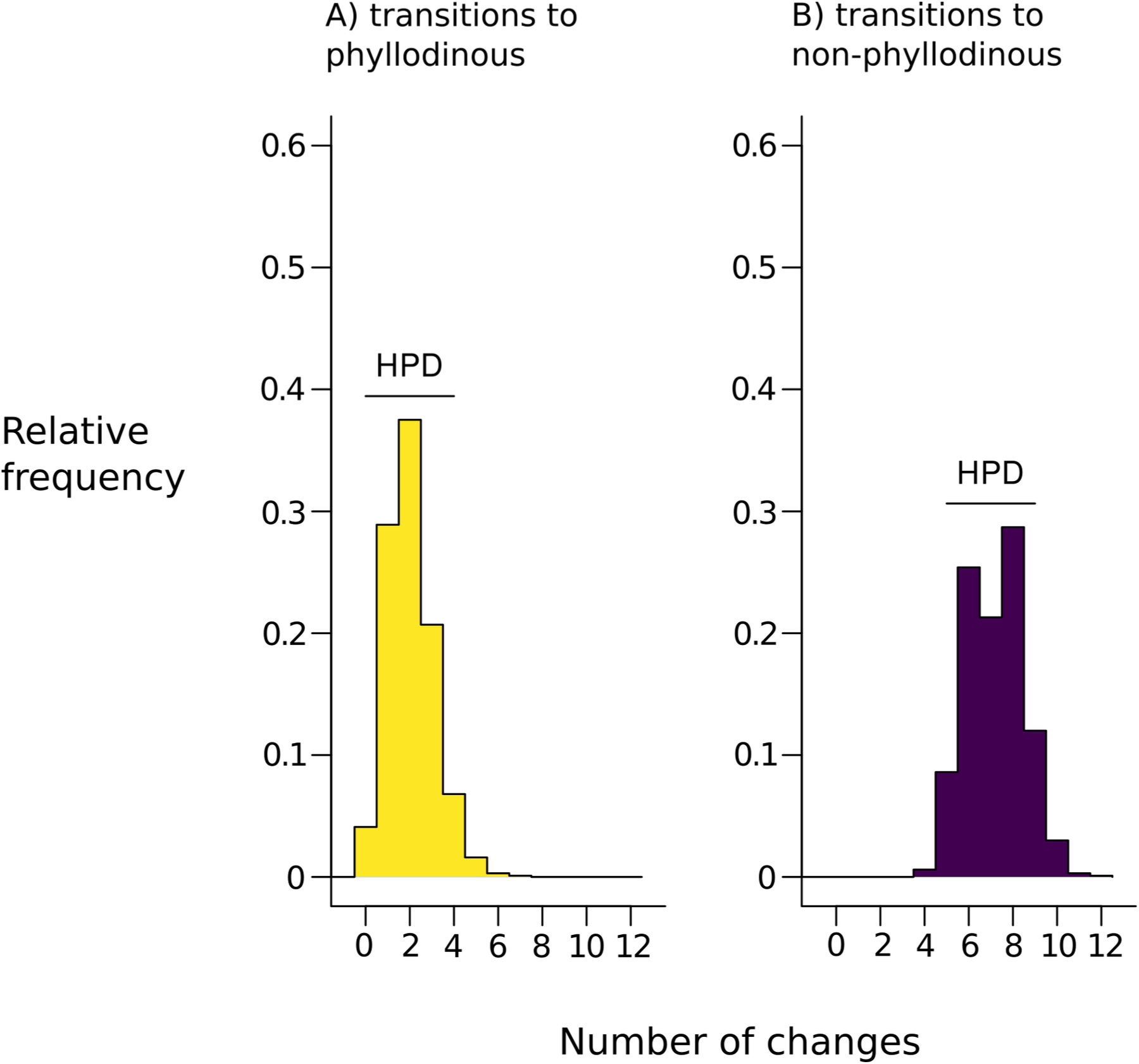
Distribution of heterochronic shifts from the 1000 simulated mappings. (A) bipinnately compound to phyllodinous and (B) phyllodinous to bipinnately compound from the ancestral character state reconstruction (Figure 2). Bayesian 95% high probability density intervals (HPD) are plotted over the distributions (from 0-4 for A, and 5-9 for B).

## TABLES

**Table S1.**
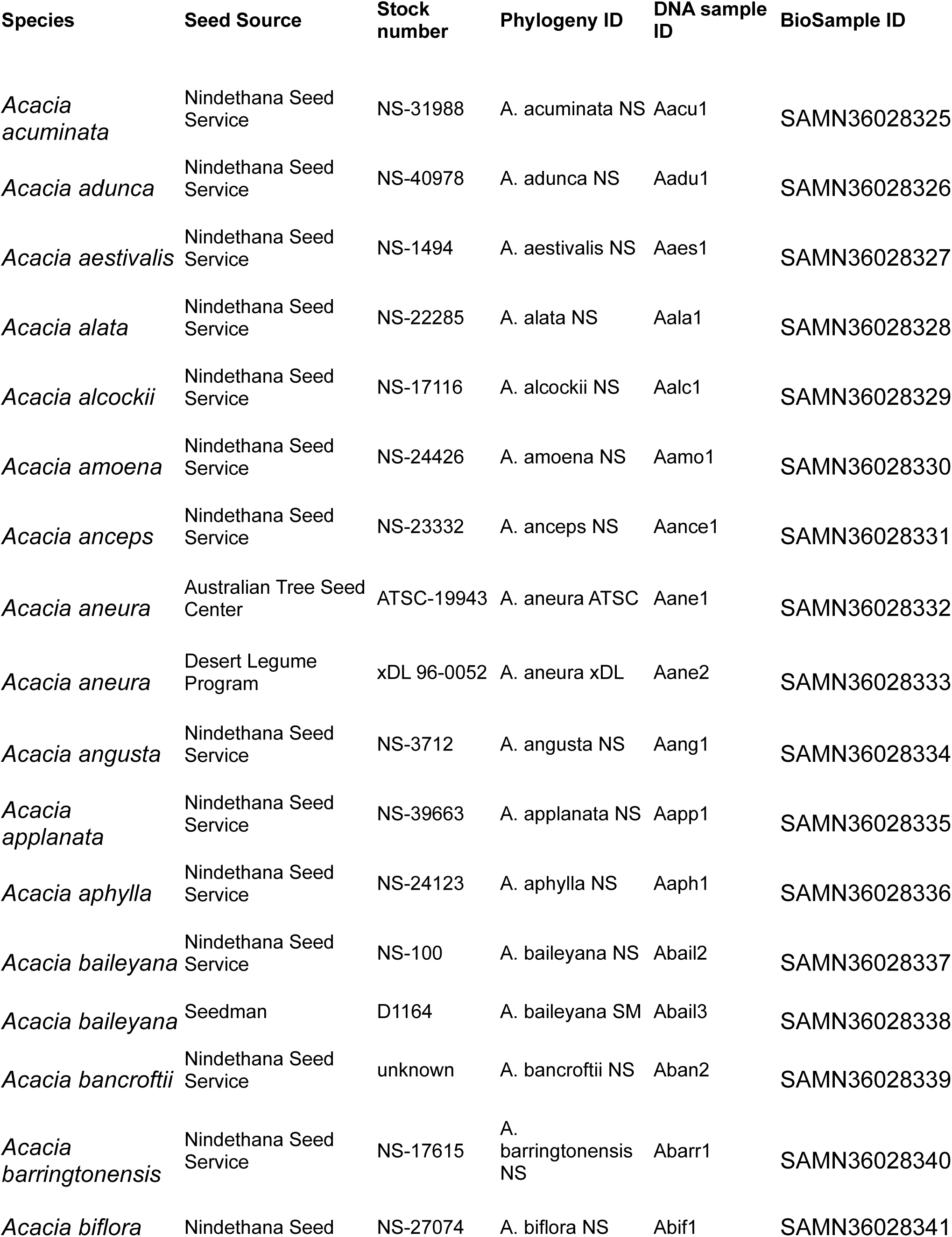
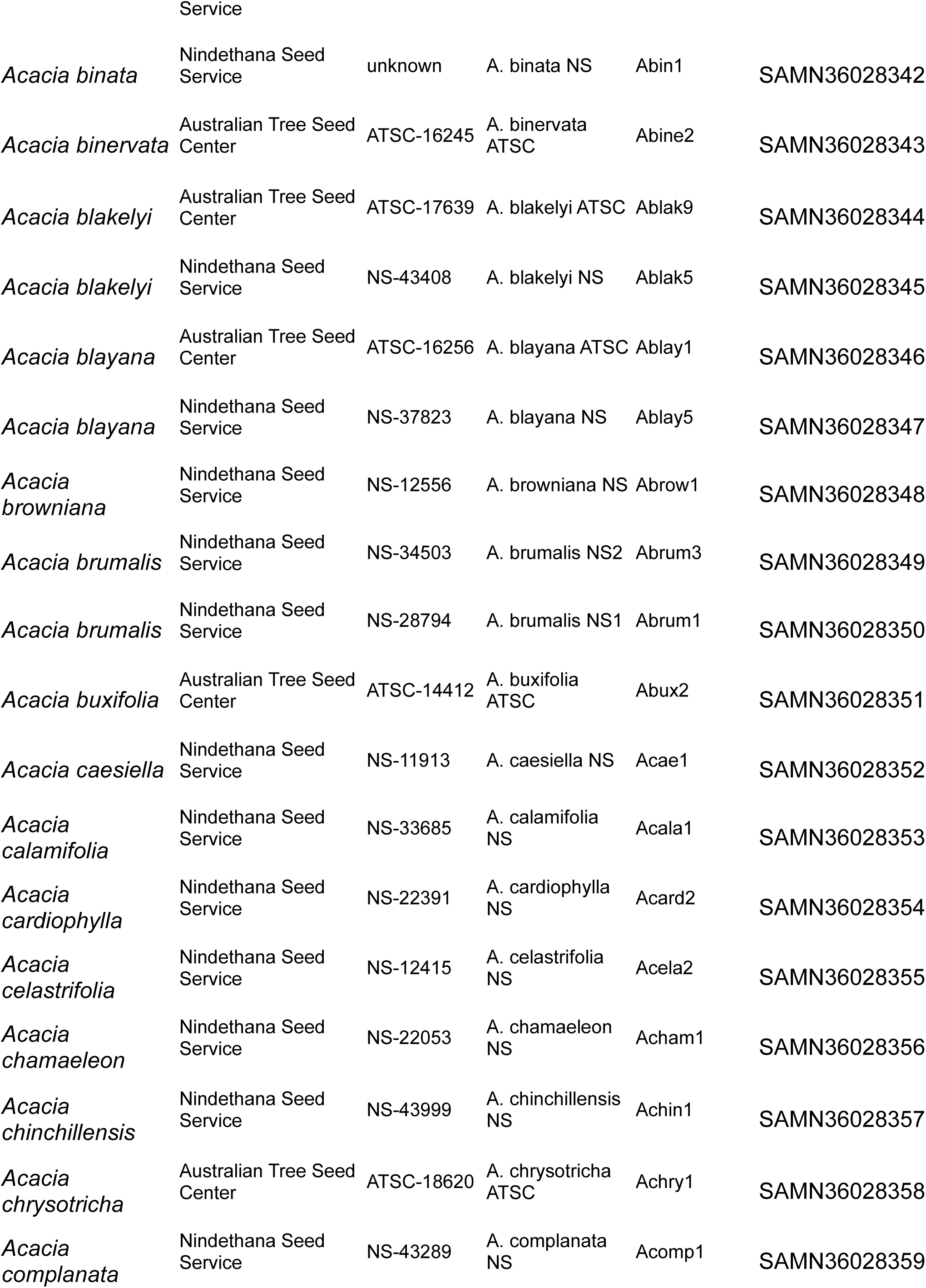
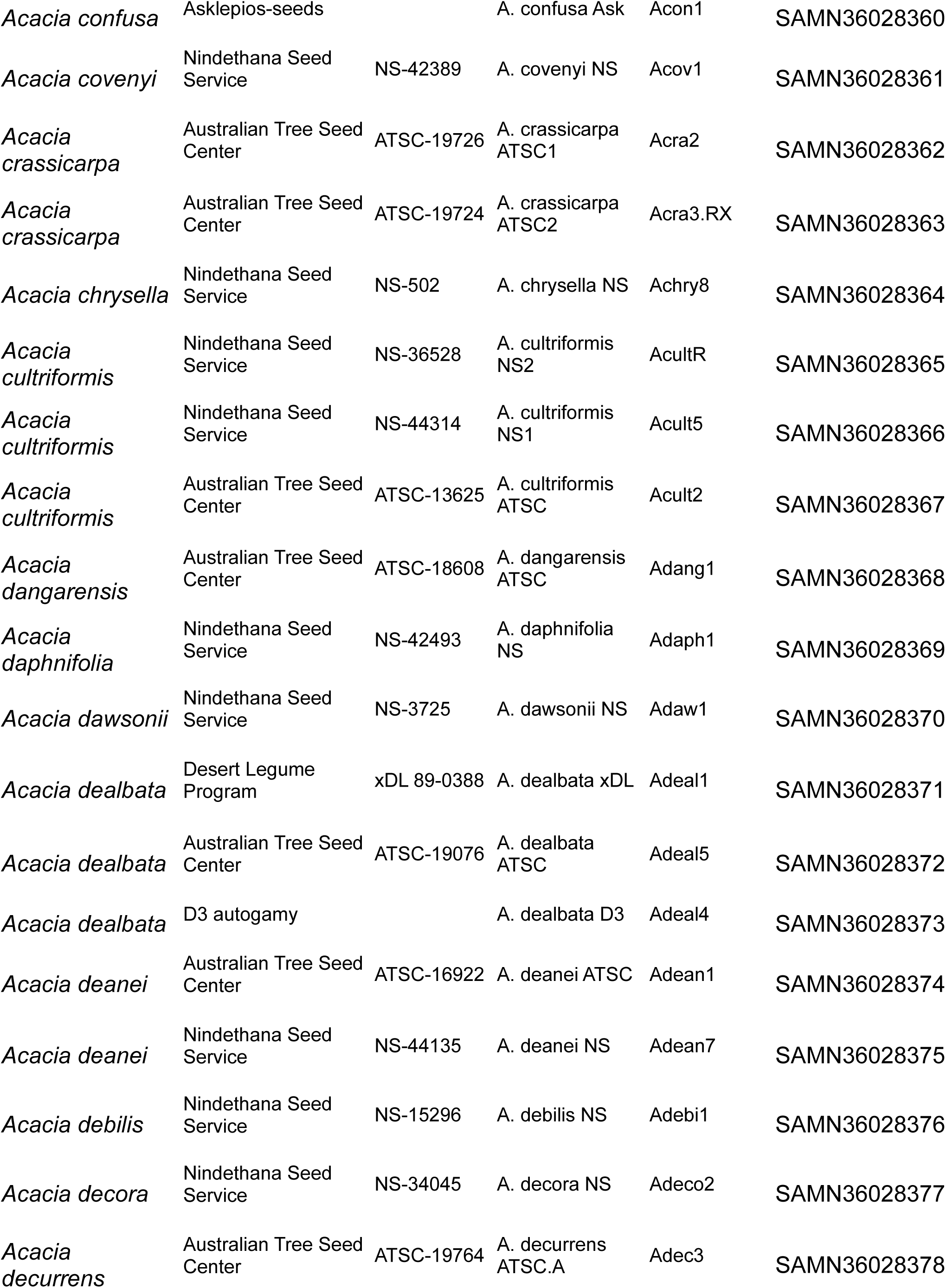
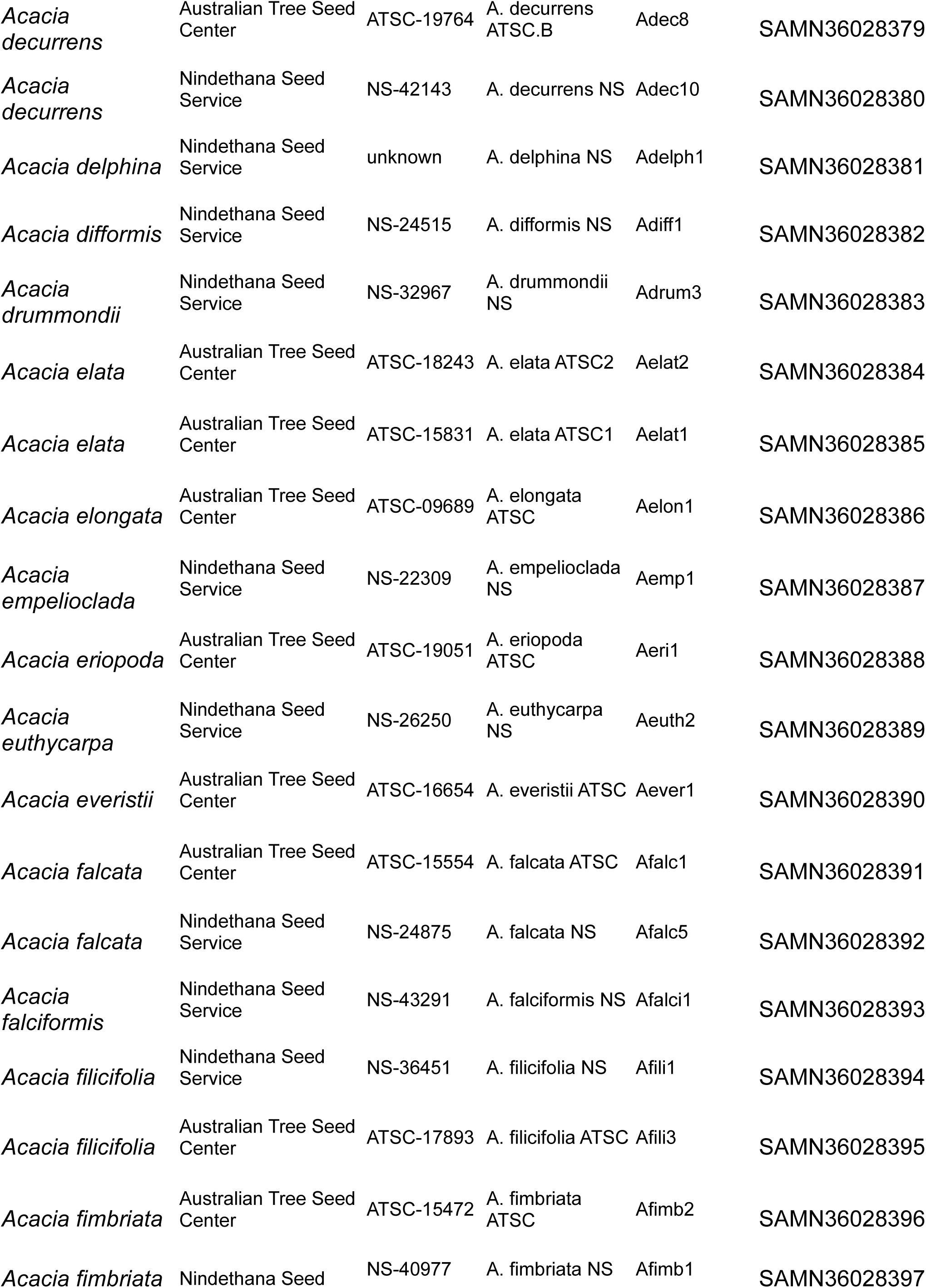
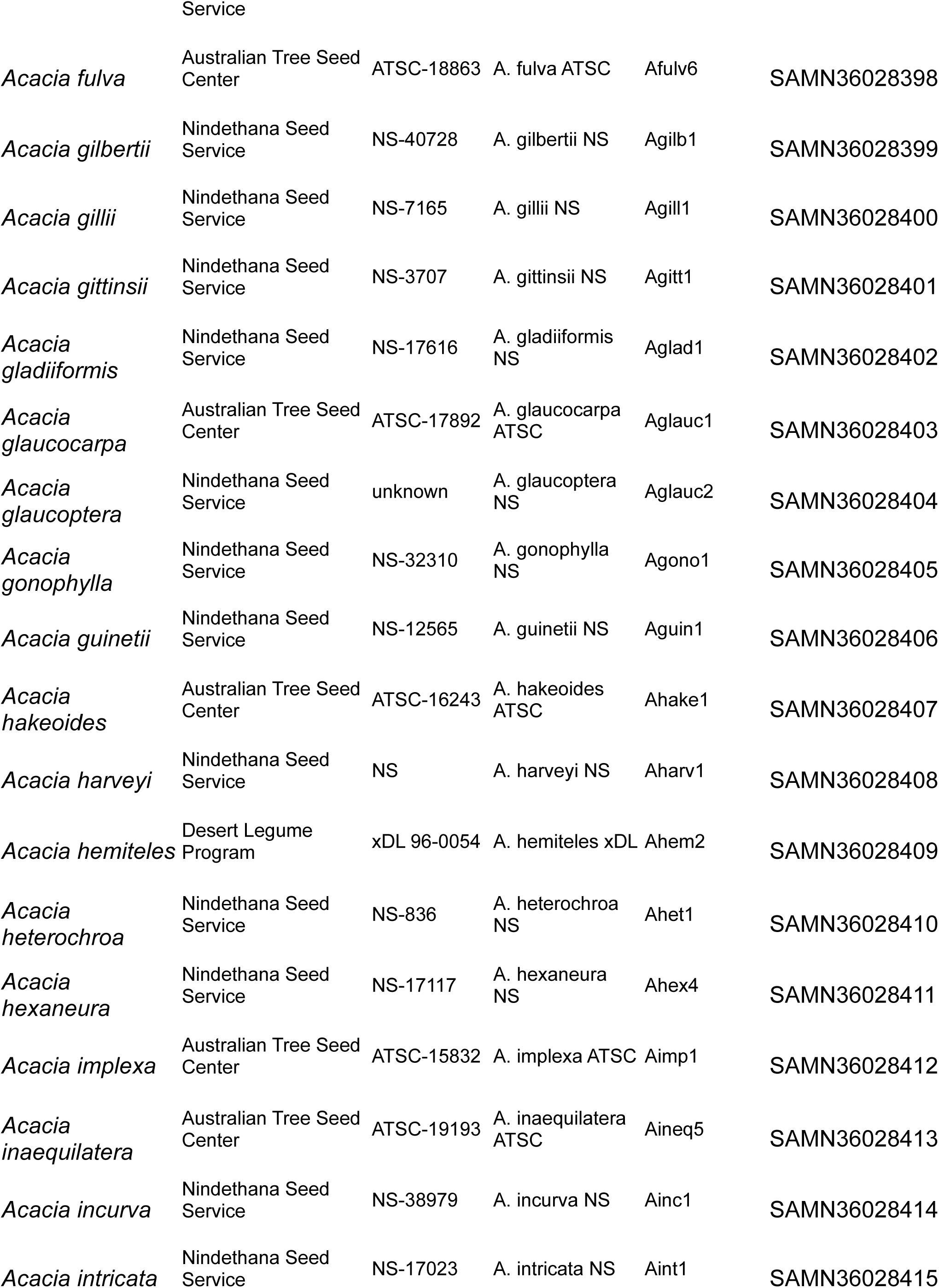
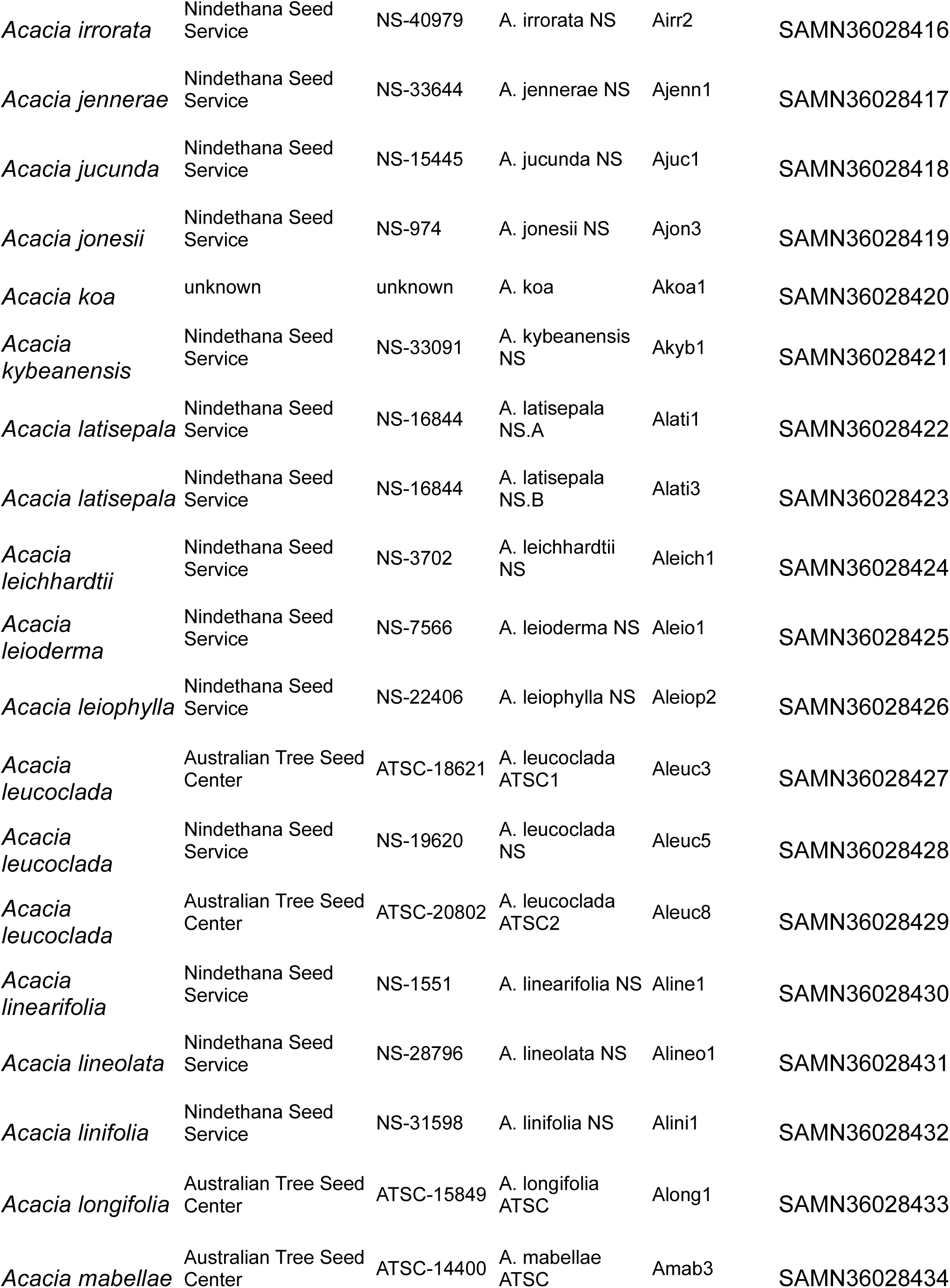
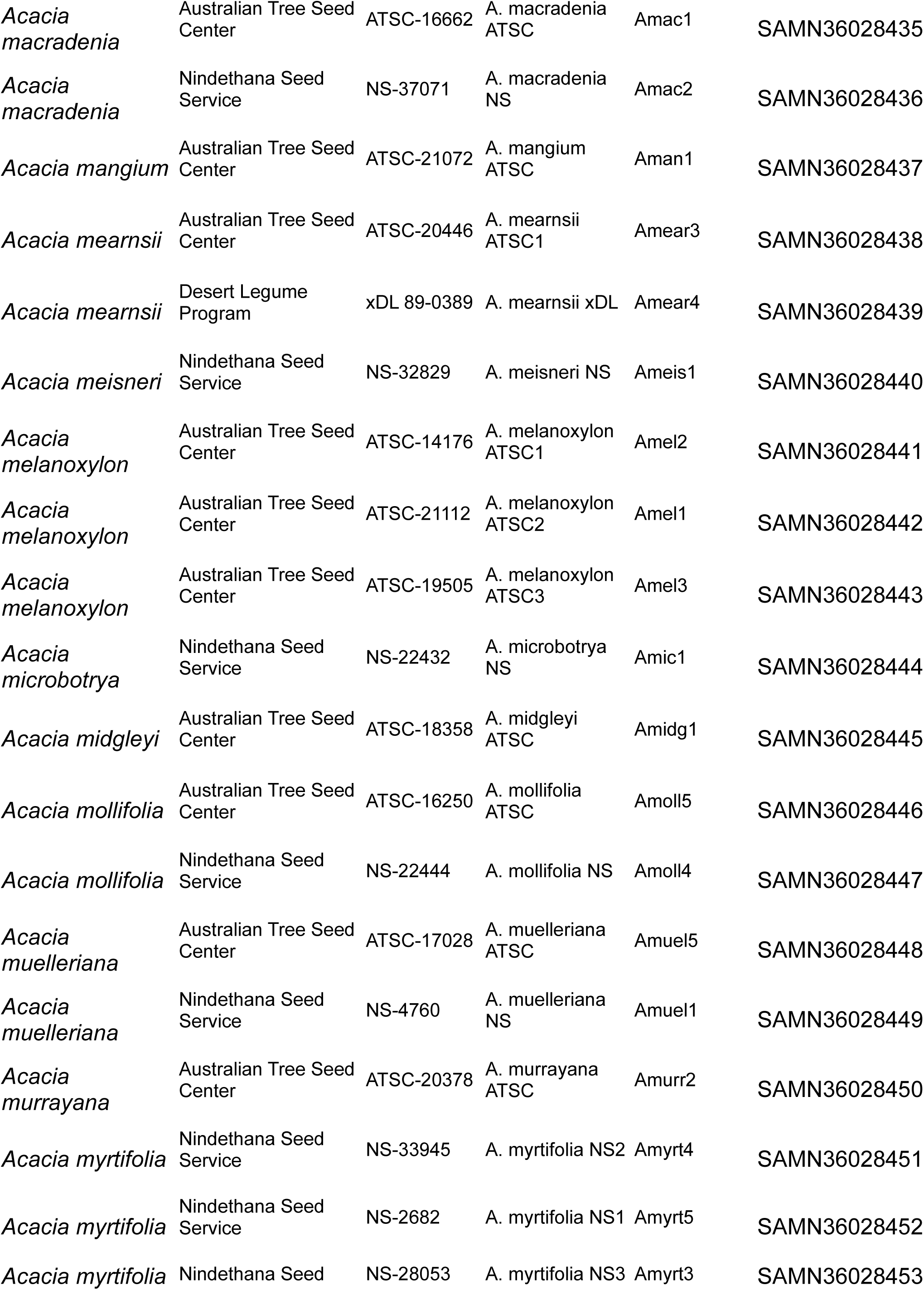
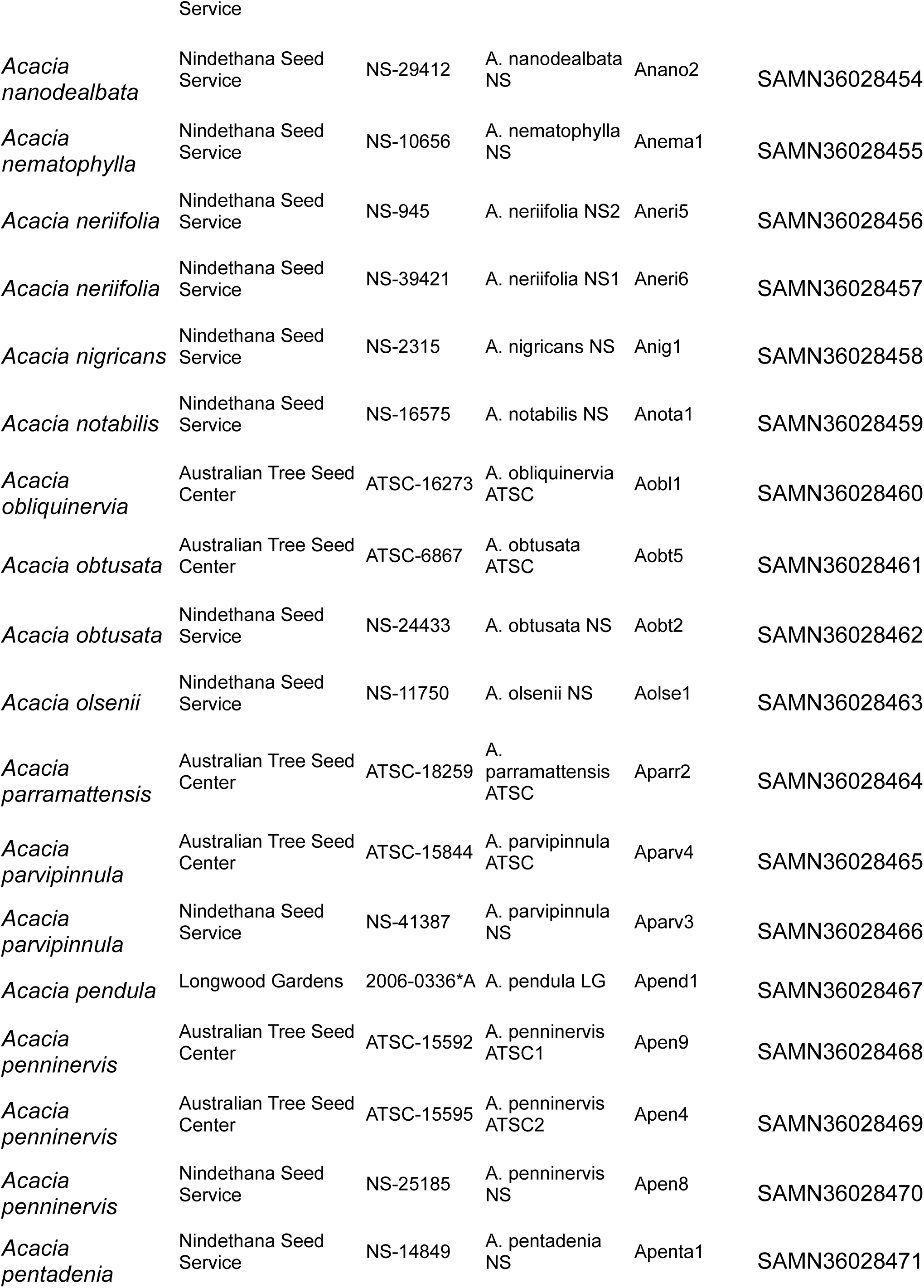
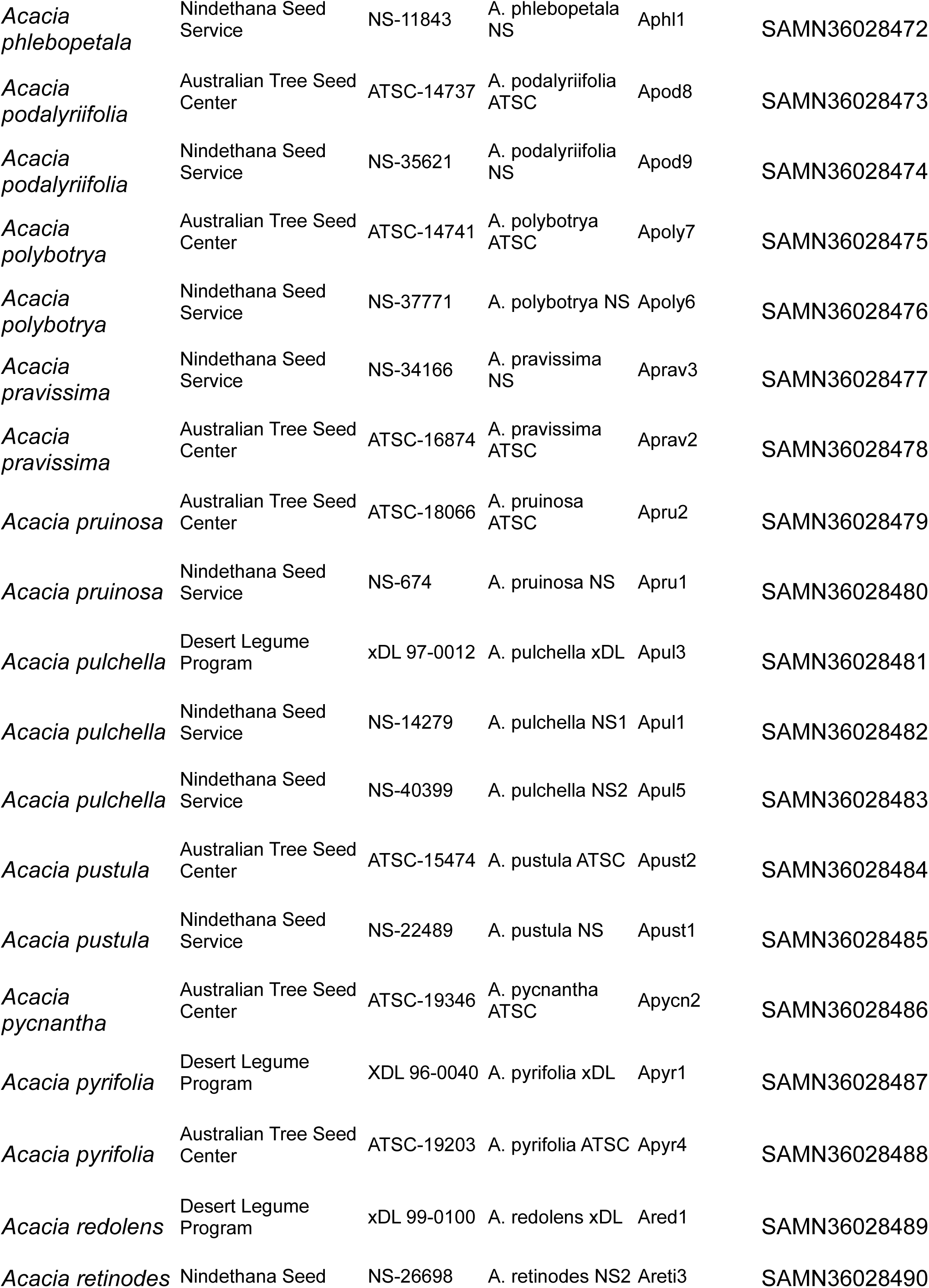
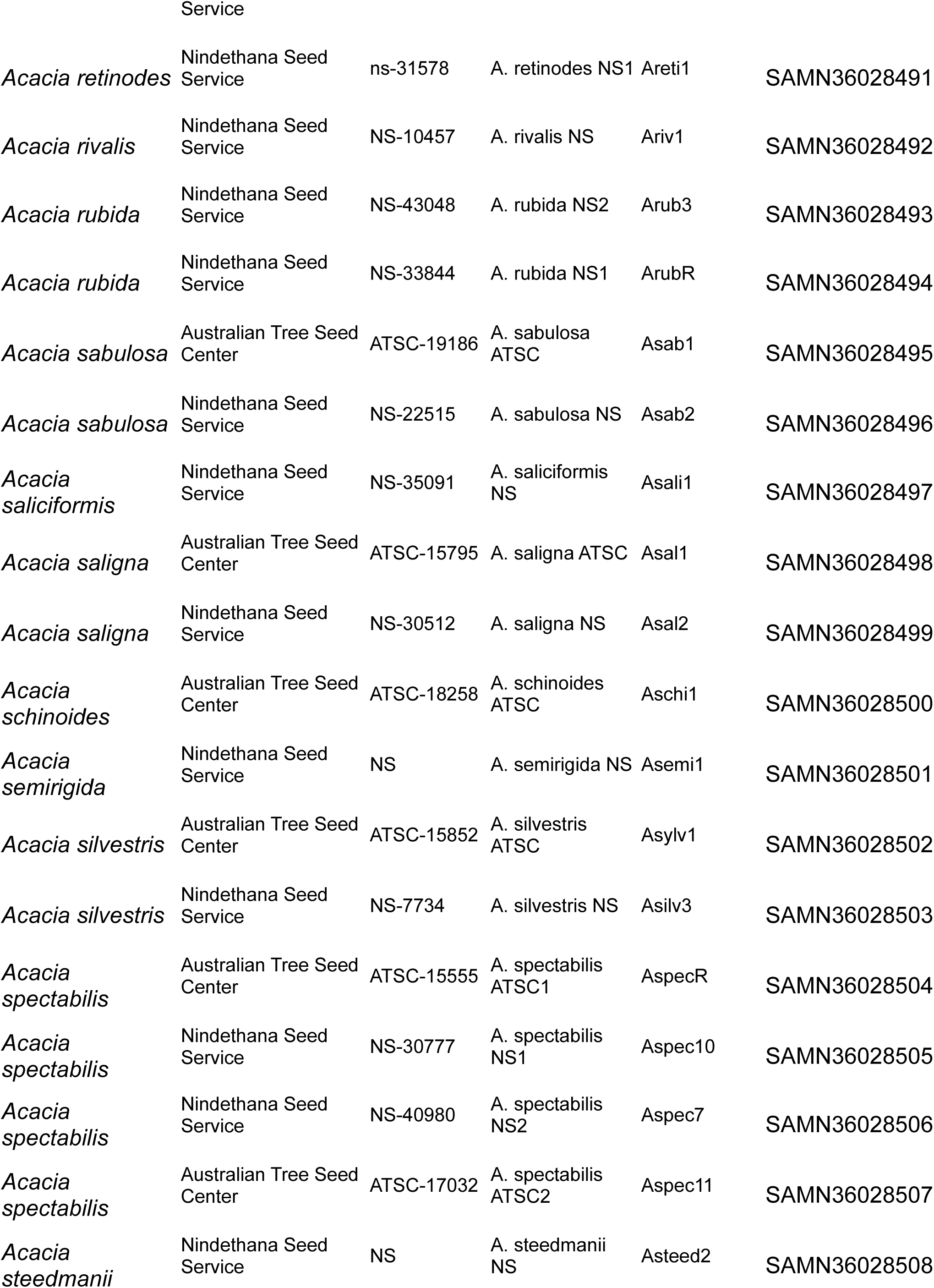
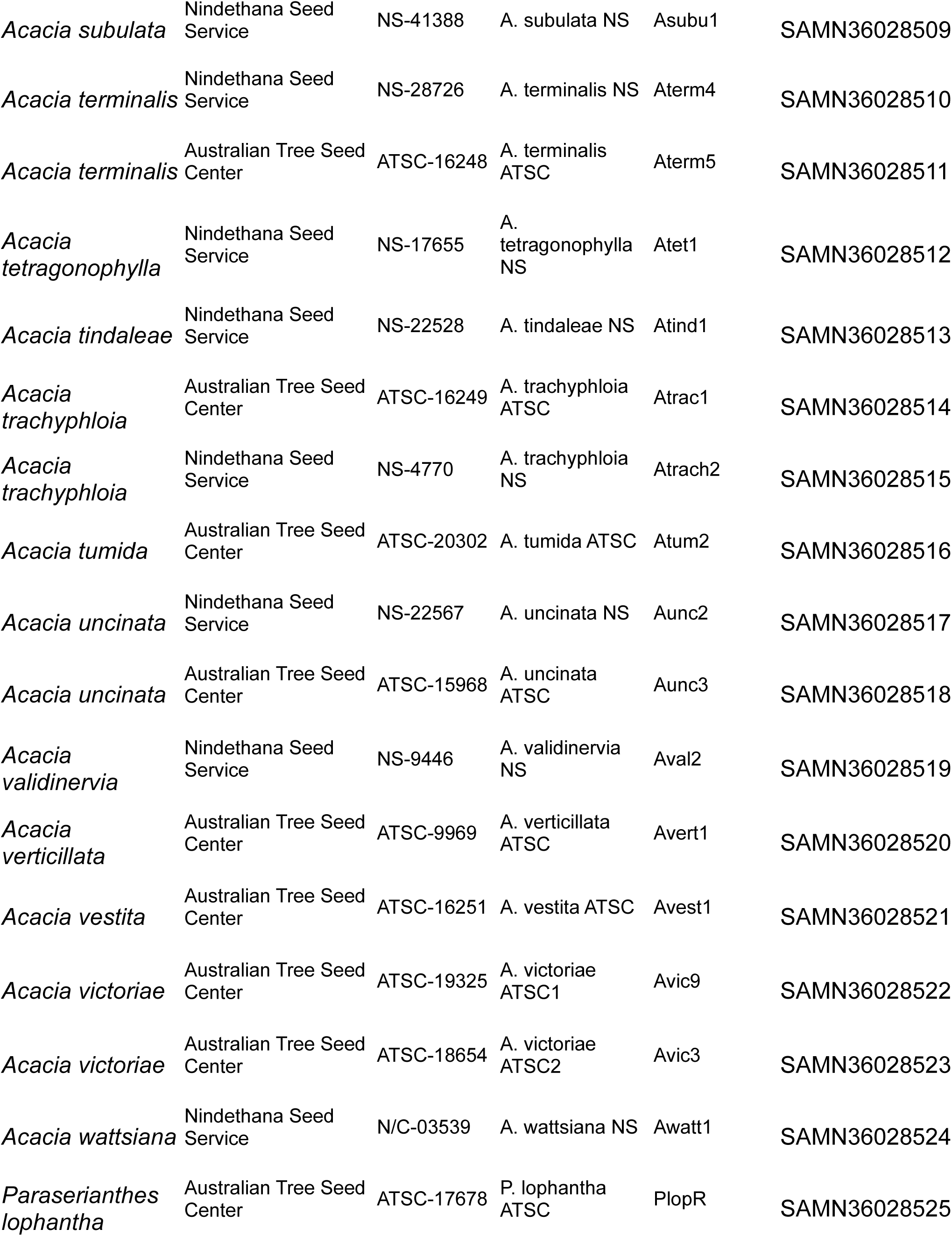
Full list of samples used for ddRAD-seq.

**Table S2.**
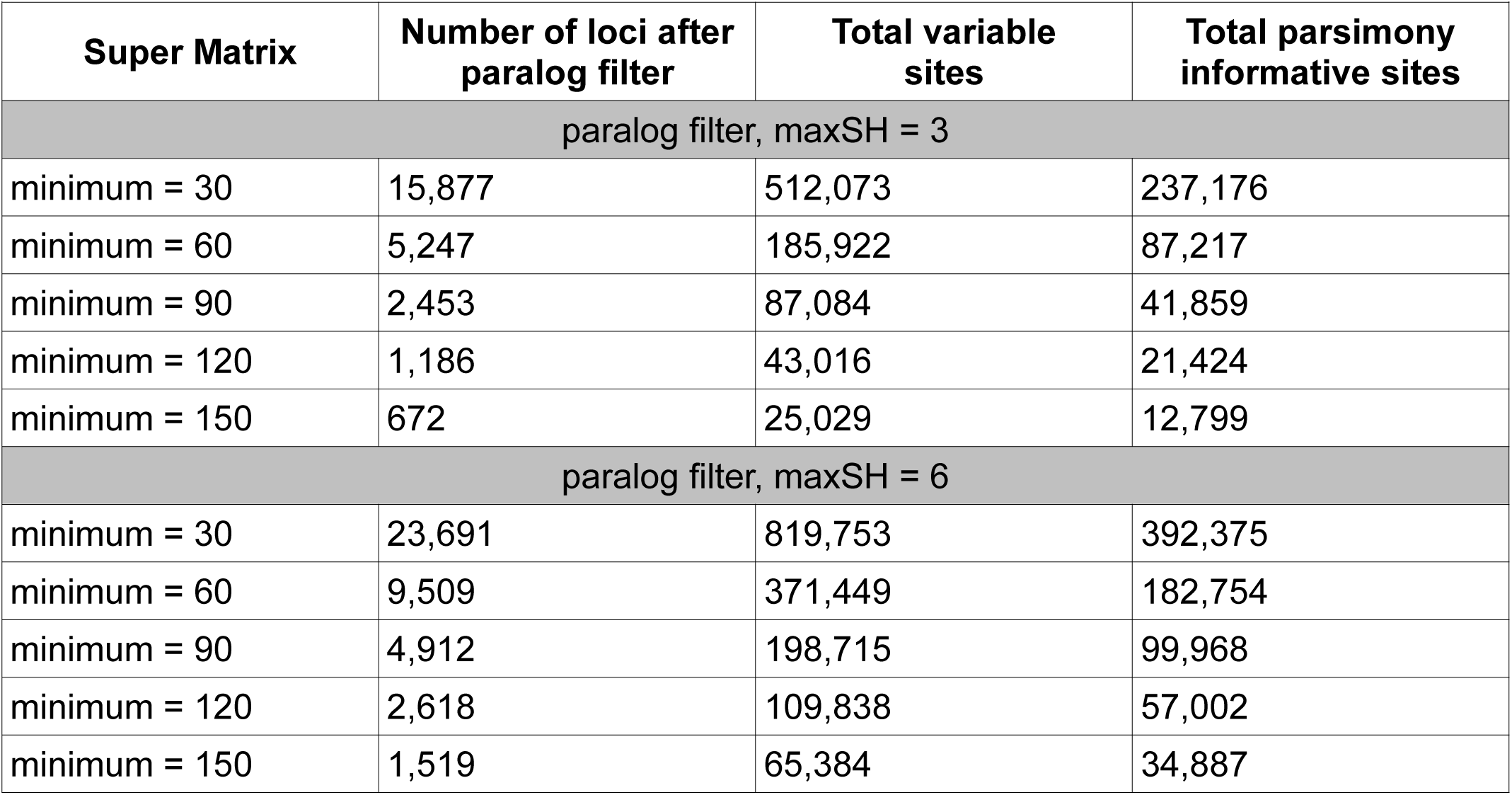
Size of supermatrices generated from the ddRAD-seq data.

**Table S3.**
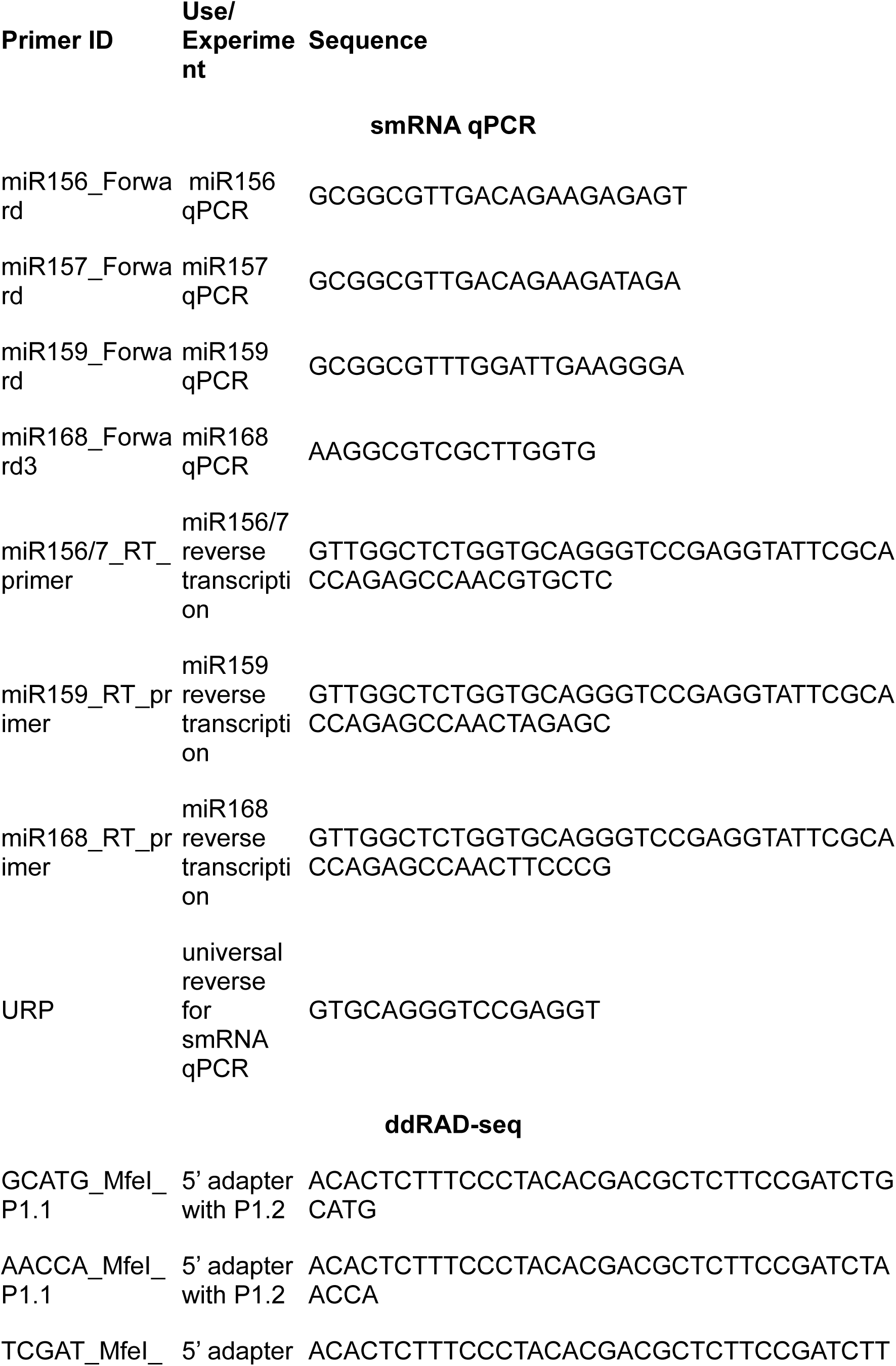
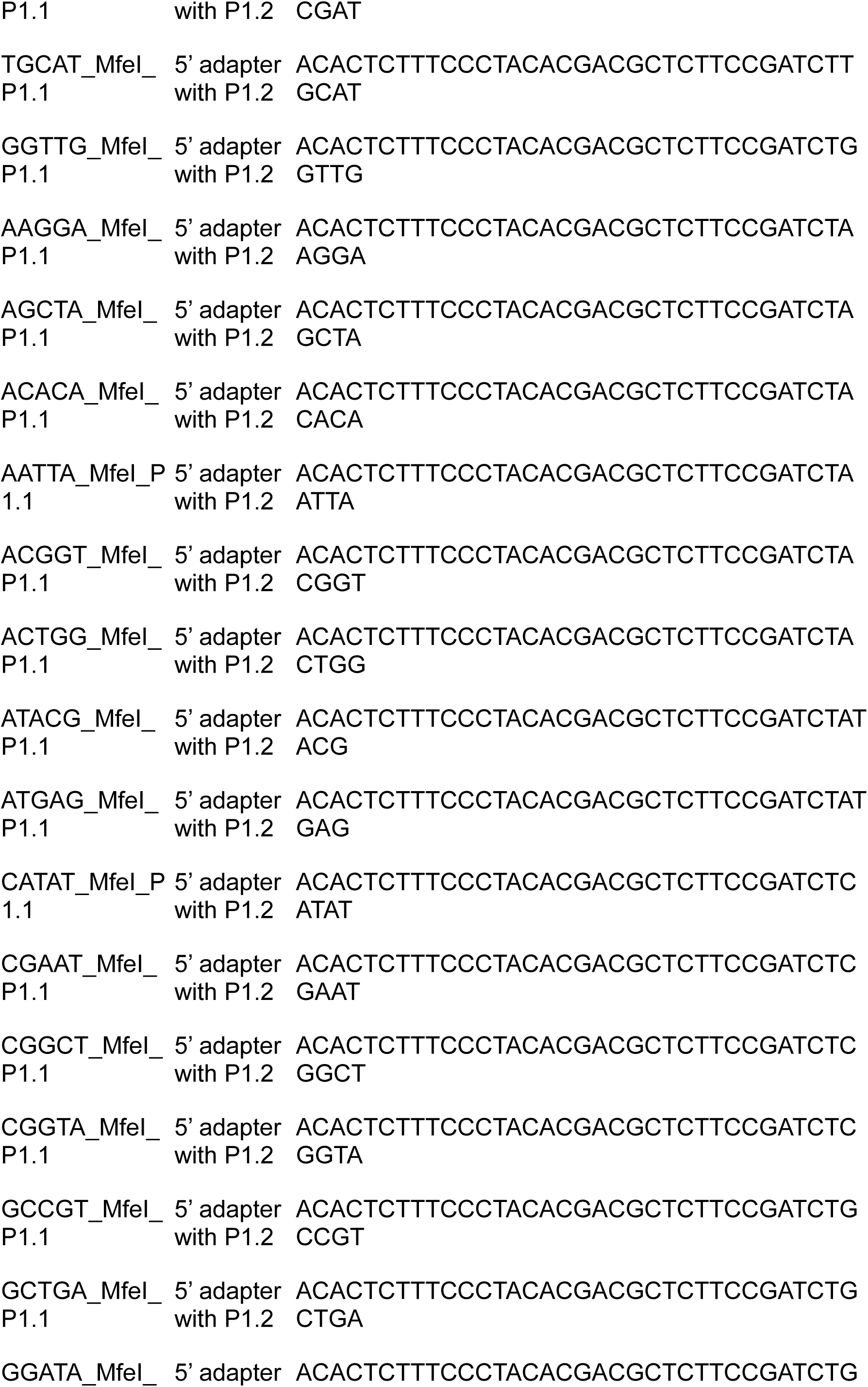
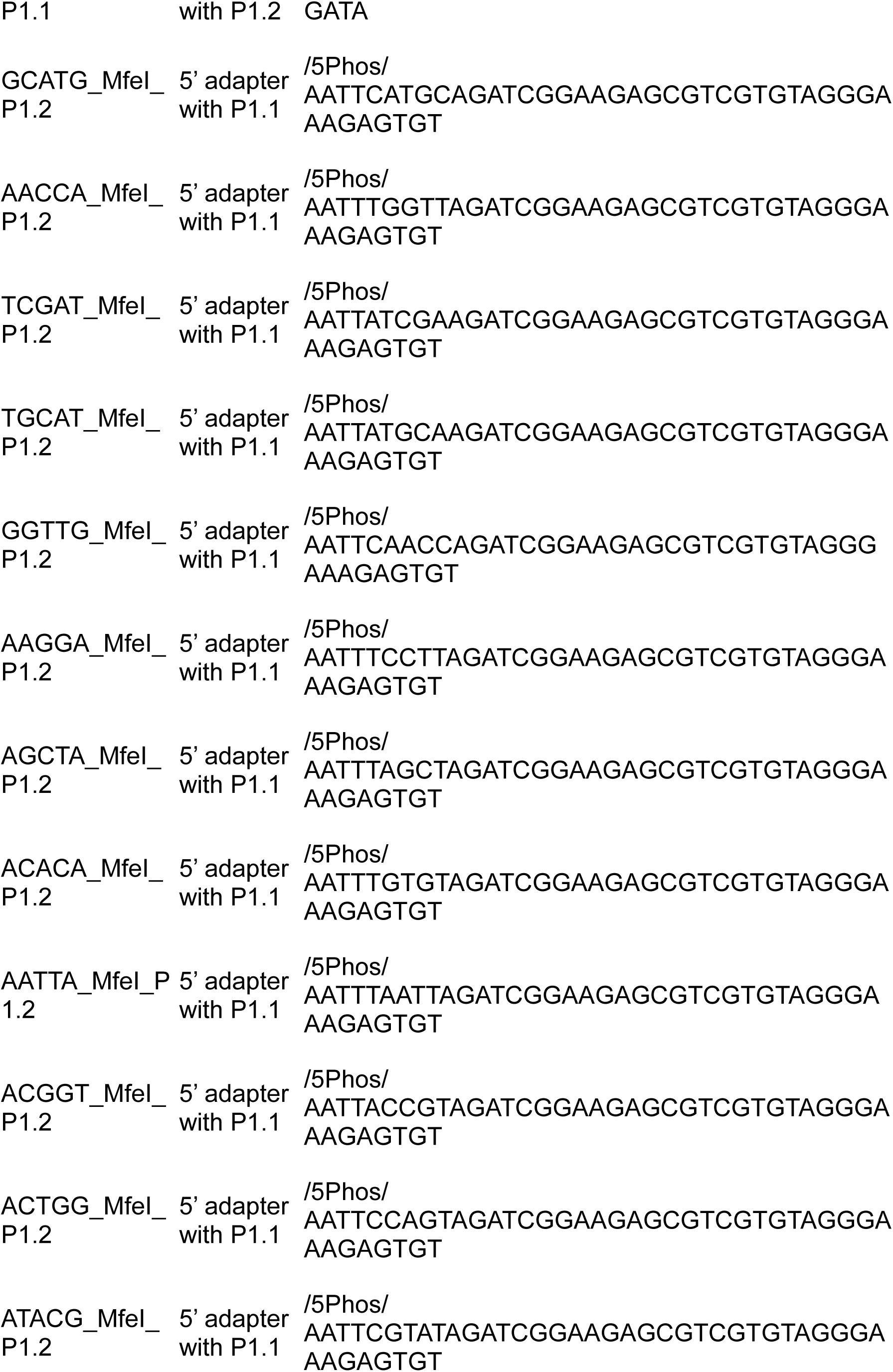
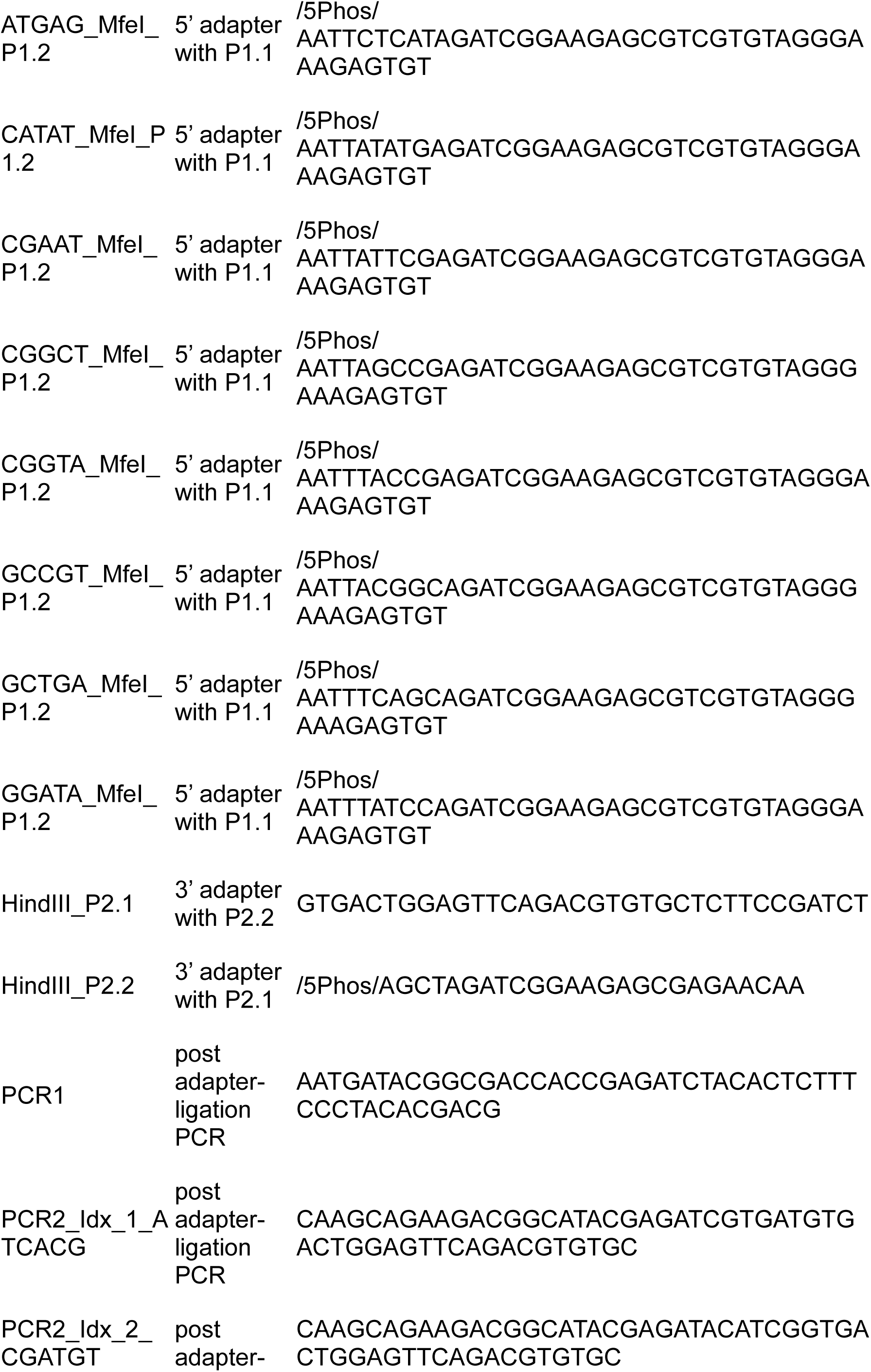
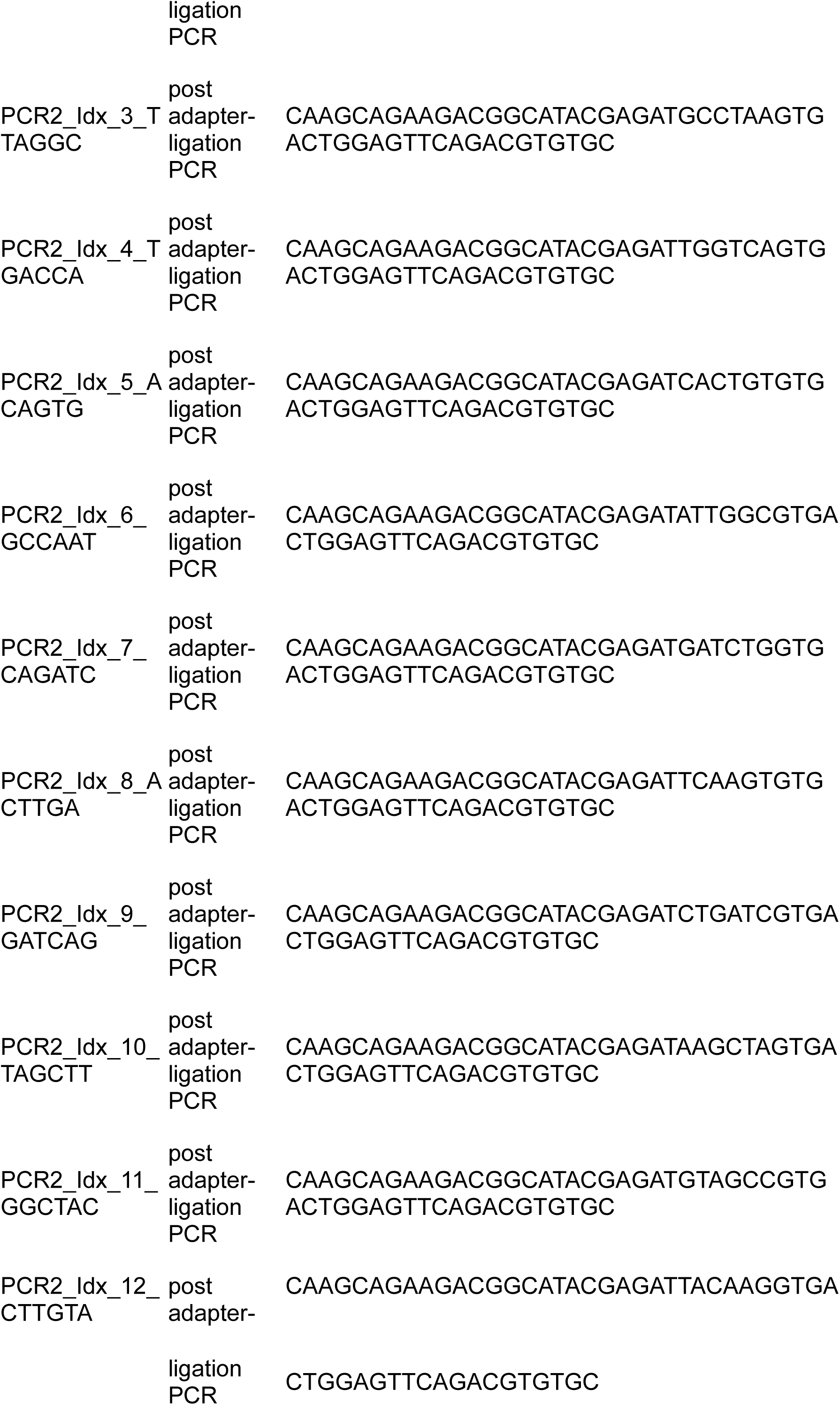
Primers used in this study.

